# Boosting class II HLA epitope presentation to T cells with endoplasmic reticulum transmembrane domain fusion proteins

**DOI:** 10.1101/2023.06.22.546056

**Authors:** Veronica Pinamonti, Ana Mellado Fuentes, Theresa Schmid, Miray Cetin, Taga Lerner, Miguel Hernandez Hernandez, John M. Lindner

## Abstract

Antigen processing and presentation are crucial for T-cell receptor engagement and potent adaptive immune responses. While establishing an antigen presentation system to identify TCR-epitope pairs from large libraries of genetically templated, ectopically expressed antigens, we observed a longitudinal suppression of antigen processing in B cells, which we call antigen processing attenuation (APA). Beyond its potential role in regulating B cell-enforced peripheral tolerance, APA represents a bottleneck to sustained antigen presentation in screening and vaccination approaches. To overcome APA we screened several pathways which might enhance processing. Notably, we identified several sequences derived from transmembrane domains (TM) of endoplasmic reticulum-resident proteins which promote sustained minigene-encoded antigen presentation. In particular, the first TM domain of *SYT6*, a protein associated with synaptic vesicles, greatly enhances processing and ameliorates APA. These findings suggest a new entry point into antigen processing pathways, enabling the discovery of novel immune epitopes and improving T-cell based immunization strategies.

## Introduction

Antigen processing and presentation utilize several complex and elaborate pathways, involving numerous enzymes and adapter proteins which ultimately enable T cells to detect pathogenic infections or genomically disruptive perturbations. The proper control of T-cell responses *in vivo* is crucial for maintaining homeostasis, protecting against infectious disease, and preventing cancer and autoimmunity. Therefore, the identification of foreign, self-, and neo-antigens presented by APCs is an essential step in the prevention, diagnosis, and therapy of such events. While high-throughput techniques for sequencing T-cell receptor repertoires are available^1^, the robust deconvolution of TCR specificities remains a technical challenge. The main obstacles to these approaches are HLA diversity^2^ (more than 12,000 known alleles) and the ability to screen a proteome-wide library simultaneously for class I-and class II-presented epitopes. Class I HLA molecules present antigens derived from cytoplasmatic proteins following proteasome-mediated degradation^3^ (for recognition by cognate cytotoxic CD8+ T cells^4^), while class II HLA molecules typically present extracellular antigens following their internalization and processing in lysosomal vesicles^5^ to CD4+ T cells^6^. This mechanistic segregation would seemingly present a significant challenge to deconvolution strategies featuring genetically encoded putative cognate antigens, particularly if class II HLA-presented targets are to be expressed intracellularly. However, the division of cytoplasmic and extracellular antigen presentation by class I and class II HLA, respectively, is not absolute, and several pathways have been identified by which intracellular epitopes can be presented by class II HLA complexes, and vice versa^7, 8^. As a key feature of a more robust functional platform, we aimed to identify and leverage one or more of these pathways to ensure the presentation of class II epitopes to CD4-restricted TCRs.

To establish our antigen-presenting system for T-cell antigen identification, known as T-FINDER (T-cell Functional Identification and (Neo)-antigen Discovery of Epitopes and Receptors, described in Schmid and Cetin, et. al., 2023 BioRxiv^9^), we exploited endogenous processing and presentation pathways to enable simultaneous class I and class II antigen presentation. Our antigen-presenting constructs and libraries are composed of synthetically assembled open reading frames (“minigenes”), the rapid design of which is partially enabled by our Fast Algorithm for Codon Optimization (FALCON), a script developed for nucleotide back-translation. One crucial benefit of our approach to antigen presentation is the natural processing of genetically encoded antigens by the intracellular machinery, which avoids potential issues and artifacts associated with forced presentation of non-physiological epitopes, variable peptide lengths, *cis*-spliced peptides^10, 11^, and haplotype-specific presentation events. While optimizing T-FINDER, we encountered a previously unexpected challenge: the longitudinal suppression of class II HLA antigen processing from longer minigene constructs. If not addressed, this hurdle would critically limit the maximum length of minigene constructs and significantly increase the size of antigen libraries used for high-throughput *de novo* ligand identification screens – rendering the evaluation of such libraries for cognate targets at sufficient depth statistically infeasible. We therefore sought to verify this issue as a *bona fide* phenomenon, then engineer solutions to overcome it. As a secondary benefit, we aimed to find solutions that would more generally enhance antigen presentation from larger minigenes, which typically do not induce as potent activation signals in reporter T cells as, for example, *in vitro*-loaded peptides or CD74 fusion constructs^12, 13^.

## Results

### Design principles for improved processing and presentation of transgenic class II HLA epitopes

To guarantee natural processing of antigens encoded by our library, we designed a lentiviral construct which allocated a 400 amino acid open reading frame (ORF) for minigene expression (“2B”, linked via a 2A viral peptide^14^ to mTagBFP for in-frame and expression validation). Here, the term ‘minigene’ describes any combination of one or more full-length proteins or fragments thereof joined by flexible peptide linkers (Supplementary Figure 1A). Full-length proteins longer than 400 amino acids were deconstructed into fragments of the specified length and overlap, then (if necessary) reassembled to generate a list of equally sized minigene sequences (Supplementary Figure 1B). We selected an overlap of 30 amino acids to avoid incorporating joints that may interrupt relevant epitopes^15^. Using this approach, we estimate that an antigen-presenting library containing 4×10^4^ elements will cover the entire human reference proteome (20,000 protein-coding genes, length distribution in Supplementary Figure 1C)^16^ at 100-fold screening depth. Notably, this approach reduces by 10-fold the number of elements required for a full-proteome screen relative to using 50 amino acid minigenes (Supplementary Figure 1D), making proteome-wide screening at sufficient depth feasible. After defining their amino acid composition, sequences are back-translated with our Fast Algorithm for Codon Optimization (FALCON), which generates nucleotide sequences that eliminate restriction enzyme recognition sites and undesired nucleotide motifs (homopolymer stretches, AT- and pyrimidine-rich stretches) while normalizing GC content (Supplementary figure 1E, 1F). Most importantly, FALCON normalizes codon usage, which depends on tissue-specific tRNA abundance^17^, and codon autocorrelation^18^ to the cell type used experimentally (in this case, EBV-immortalized B cells), potentially increasing mRNA stability and/or the efficiency of protein translation^19, 20^. Relative to other available tools, FALCON considers more parameters while more quickly generating sequences, enabling the rapid backtranslation of large protein libraries^21, 22^ (Supplementary Table 1).

Our antigen (library) presenting system aims to simultaneously process and present class II and class I epitopes from larger ORFs encoded by a single transgenic construct. To test the suitability of our design for this purpose, we selected as class I presentation controls the TCR DMF5^23^ and the MLANA^wt^ gene, which encodes the MART-1 epitope, and MLANA^A27L^ for the MART-1^A27L^ high-affinity epitope^24^. For class II presentation, we selected the tetanus toxoid-reactive TCR TT2 (Supplementary Table 2) and its cognate epitope-containing fragment from the *C. tetani* tetanospasmin protein (TetX 1 – 399). Both minigenes were cloned into the lentiviral construct and used to transduce BOLETH cells (an EBV-immortalized B-cell line)^25^, while TCRs were expressed in the Jurkat-based TCR reporter PNJE3 cell line^9^, which expresses GFP upon T-cell activation. In both cases, strong TCR activation was detected, indicating that both class I and class II presentation are possible in EBV-immortalized B cells when transduced with lentiviral particles encoding 400 AA minigenes (Figure 1A). We next tested whether the primary minigene sequence (permutations of the natural protein compositions) might affect T-cell activation potential. PNJE3 cells expressing the DMF5 TCR were co-cultured with B cells expressing the MLANA natural ORF, or a 400-amino acid minigene containing the MART-1 peptide epitope embedded among irrelevant sequences. Additionally, we tested a pair of minigenes containing the cognate epitope for another tetanus toxoid-reactive TCR (TT7) at either the 5’-or 3’-end of a minigene. Strong T-cell activation was detected in both cases, confirming that the minigene composition does not generally affect epitope presentation (Figure 1B, Supplementary figure 1G).

**Figure 1.**
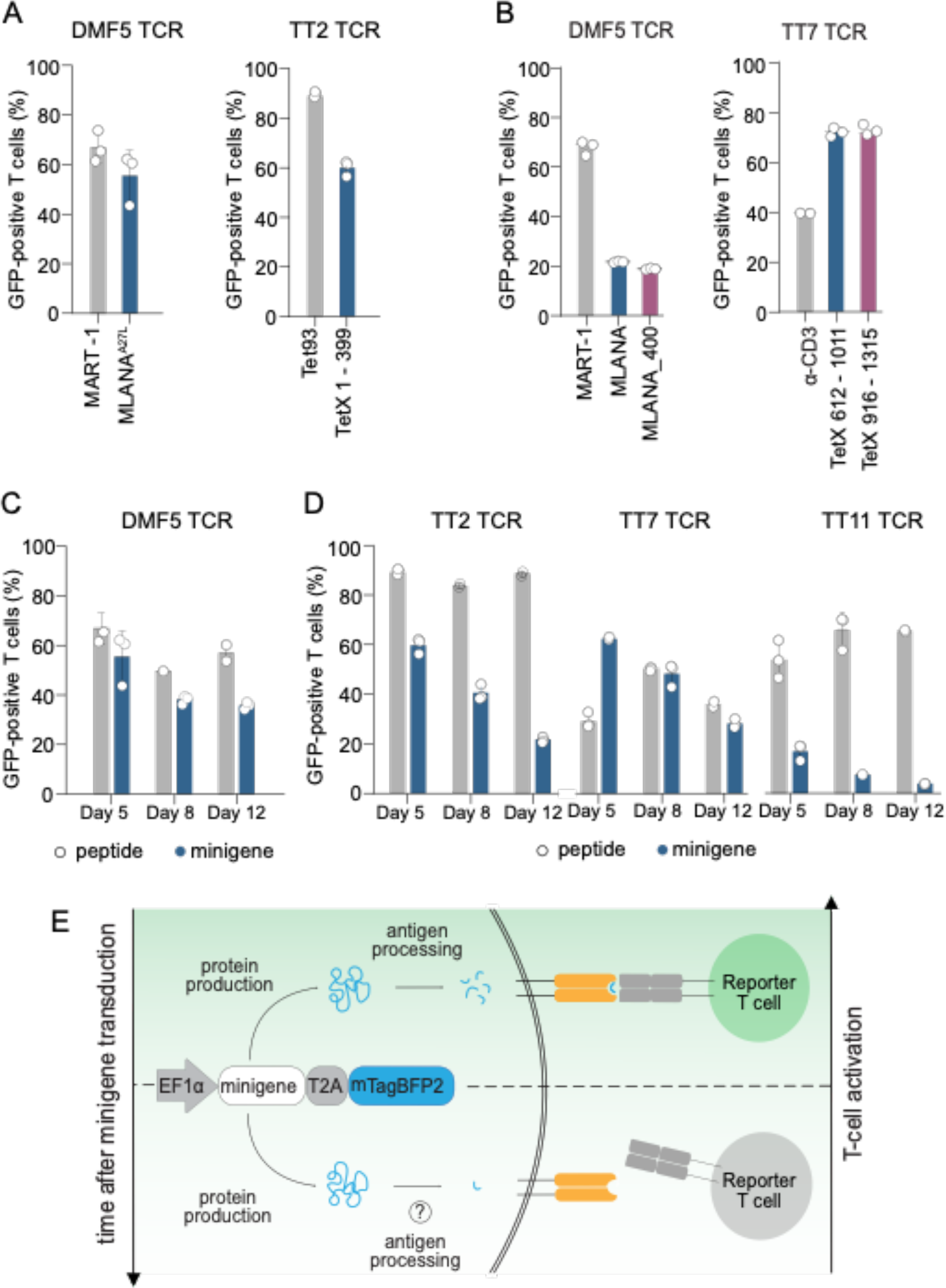
Transgenically introduced minigenes induce class I and class II presentation. (A) Co-culture of positive control ligand-containing minigenes (blue bars) for antigen presentation using minigene-expressing BOLETH cells and PNJE3 reporter cells transduced with the cognate TCR. For class I, a MLANA^A27L^ minigene and the DMF5 TCR were used; or class II, a TetX 1 – 399 minigene was paired with the TT2 TCR. Positive controls (gray bars) used BOLETH cells loaded *in vitro* with 10 μM MART-1 peptide or with 1 μM Tet93 peptide. Results are shown as the percentage of GFP positive T cells. (C) Left, co-culture of BOLETH cells pulsed with 10 μM MART-1 peptide or transduced either with MLANA^wt^ full-length ORF or MLANA_400 minigenes and co-cultured with reporter cells expressing DMF5. Right, co-culture of BOLETH transduced with TetX 916 – 1315 (epitope N-term) or TetX 612 – 1011 (epitope C-term) with reporter T cells expressing the TT7 TCR. As a positive control, reporter cells were stimulated with 2.5 μg/ml α-CD3. Results are shown as percentage of GFP positive T cells. (D) Longitudinal co-culture experiment performed at days 5, 8, and 12 post-transduction using the MLANA^A27L^ minigene or 10 μM MART-1 pulsed peptide and the DMF5 TCR. (E) Longitudinal co-culture experiment for class II HLA-presented epitopes using the TetX 1 – 399 minigene and TT2 TCR, or TetX 612 – 1011 with the TT11 and TT7 TCRs. Positive controls (gray bars) were 1 μM Tet93 peptide (TT2), 1 μM Tet614 peptide (TT11), or 2.5 μg/ml α-CD3 (TT7). (E) Schematic representation of a T-cell antigen-presenting system and antigen processing attenuation (APA). Minigenes linked via a T2A peptide to mTagBFP2 are processed by cellular machinery and presented by HLA molecules, leading to reporter T cell activation upon cognate interaction (top). Several days after transduction, minigene processing decreases, leading to poor or undetectable T cell activation (bottom).

The co-culture experiments were repeated longitudinally to investigate the temporal stability of antigen presentation. The class I HLA-presented MLANA^A27L^ minigene activated cognate T cells regardless whether the co-culture was performed 5, 8, or 12 days post-transduction (Figure 1C). However, T-cell activation considerably decreased over time following B-LCL transgenesis with the class II-presented antigen (Figure 1D). To validate this result we included two additional class II-restricted TCRs, TT7 and TT11 (with TetX epitopes distinct from TT2 and each other), which also exhibited reduced T-cell activation over time. To understand this decay in minigene-encoded epitope presentation, we performed a series of experiments to dissect the processes involved using TetX 1 – 399 and TT2 congate TCR. First, we confirmed that the mTagBFP2 expression levels of minigene-transduced B cells remained constant over time, excluding the possibilities of transduction bias and transgene silencing (Supplementary Figure 1H). Next, we tested whether B-LCLs stopped presenting epitopes in response to lentiviral transduction with a co-culture experiment using freshly peptide-pulsed BOLETH cells or re-pulsed minigene-transduced BOLETH (Supplementary Figure 1I). Both conditions led to strong T-cell activation, indicating that the ability of the cell to present loaded peptides is not affected by sustained ectopic expression of that (or any other lentivirally introduced) epitope. These results exclude transgenesis and presentation as potential causes of this phenomenon, leading us to conclude that antigen processing pathways play a key role in this process, which we now call “antigen processing attenuation” (APA, Figure 1E).

### Identification of presentation-boosting natural and synthetic transmembrane domains

APA represents both an intriguing immunological process and a technical hurdle to be overcome when ectopically expressing antigens for processing and presentation to T cells. Efforts to both characterize and overcome APA (Supplementary Figure 2A), including early and late lysosomal targeting^26–28^, were unsuccessful (Supplementary Figure 2B). Intervening at later stages in antigen processing using CD74 fusion proteins as previously published^12, 13^ did overcome APA, supporting our theory that the loss of antigen presentation involves the class II processing compartment (Supplementary Figure 2C). However, epitope presentation with this approach was limited to approximately 100 amino acid-long minigenes (Supplementary Figure 2D), making this construct unsuitable for our screening platform. Unexpectedly, targeting antigens to the endoplasmic reticulum using the first transmembrane domain from the B-cell chaperone Bap31^29, 30^ (BT1) N-terminally fused to cognate epitope-containing minigenes activated reporter cells strongly (similar to CD74 fusion) and stably over time using a 100 amino acid minigene for both class I (Figure 2A) and class II (Figure 2B) antigens. When using minigenes of increasing length, BT1 outperformed the 2B construct but its efficiency was still not optimal at 400 amino acids (Figure 2C). This size limit appears to be sequence-specific: when testing BT1 with additional TCR:minigene pairs, we observed strong T-cell activation at the final time point (Figure 2D). These results clearly indicate that fusing longer, antigen-containing ORFs to the first transmembrane domain of Bap31 induces strong T-cell activation and overcomes APA for at least a subset of transgenically expressed targets.

**Figure 2.**
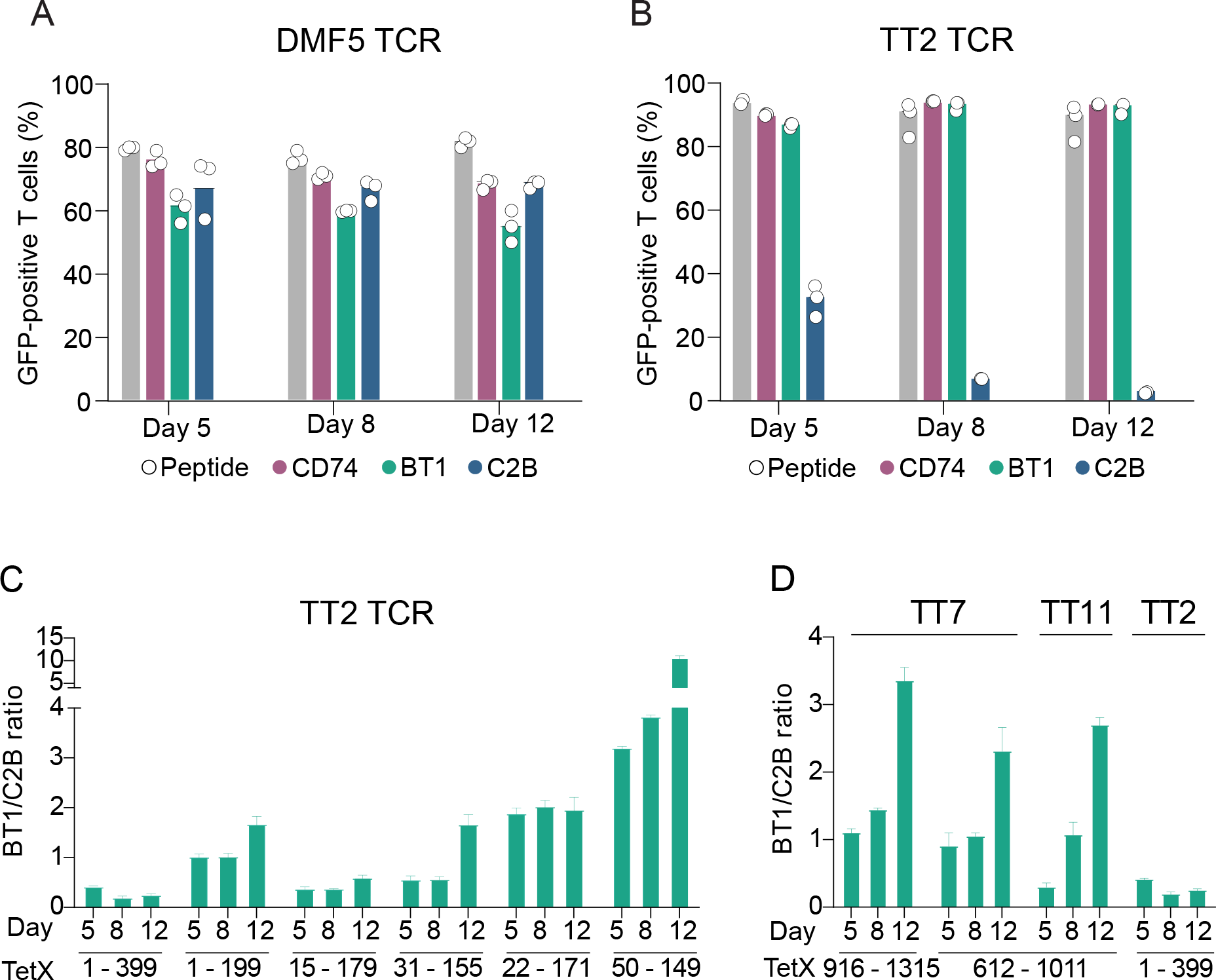
Bap31-TM1 (BT1) improves class II antigen presentation. (A) Flow cytometric analysis at day 5, day 8, and day 12 post-transduction with the MLANA^A27L^ minigene cloned in the 2B, CD74, and BT1 backbones. The experiment was performed with PNJE3 reporter cells expressing the DMF5 TCR, and 10 μM MART-1 peptide-pulsed BOLETH cells were used as an activation control (gray bars). (B) Results of co-cultures at day 5, day 8, and day 12 post-transduction with the TetX 50 - 149 minigene in the 2B, CD74, or BT1 constructs co-cultured with PNJE3 reporter cells expressing the TT2 TCR. 1 μM of Tet93 peptide (gray bars) was used to control for cognate interaction. (C) BT1 construct validation using TetX 1 - 399, TetX 1 - 199, TextX 15 −179,TetX 31 - 155, TetX 22 – 171, and TetX 50 - 149. The experiment was performed with PNJE3 reporter cells expressing the TT2 TCR. Co-culture results at day 5, day 8, and day 12 are expressed as the ratio between T-cell activation induced with the BT1 construct and the 2B construct. (D) Validation of the BT1 construct using the TetX 916 – 1315, TetX 612 – 1011, and TetX 1 - 399 minigenes with their respective cognate TCRs TT7 (two epitope-containing minigenes), TT11, and TT2. The co-culture results at day 5, day 8, and day 12 are expressed as the ratio between T-cell activation induced with the BT1 construct over the 2B construct.

The success of the BT1 fusion focused our attention on ER-resident proteins and their TM-spanning sequences for targeting minigenes to the antigen processing and presentation pathways. We randomly selected 800 sequences with a length of 21 AA from transmembrane domain sequences of ER-resident proteins. Accounting for the amino acid composition of the selected sequences, we generated an additional 400 synthetic sequences. From this library, 1152 TM domains were individually tested in a functional plate-based assay. Minigene-transduced BOLETH cells were co-cultured with reporter cells expressing the cognate TT7 TCR, and T-cell activation was measured following co-cultures at day 5 and day 12 post-transduction. We identified 18 transmembrane sequences which reproducibly induced T-cell activation across a pair of technical replicates (Supplementary Figure 3), indicating that roughly 1.5% of the library sequences do promote class II presentation (Figure 3A). ST1 (SYT6, TM1), ET1 (ESYT1, TM1), DT3 (DERL3, TM3), DA3 (DAD1, TM3), and two synthetic domains (Syn108 and Syn289) induced T-cell activation at least as strongly and stably as the BT1 fusion. These sequences were selected for further testing with 400 AA minigenes containing the epitopes of the TT11 and TT2 TCRs. For TT11, the new sequences clearly improved T-cell activation even at early time points, and delayed APA. Importantly, for the TT2 TCR, the new sequences induced strong T-cell activation with the 400 AA minigene, greatly outperforming the BT1 construct and overcoming APA (Figure 3B). A Multiple EM for Motif Elicitation (MEME-sorted)^31^ analysis of these sequences identified the common motif ‘VFLFLWK’ from the best-performing ST1 sequence, which is partially conserved in the other sequences (Figure 3C), which we call processing enhancing sequences (PES).

**Figure 3.**
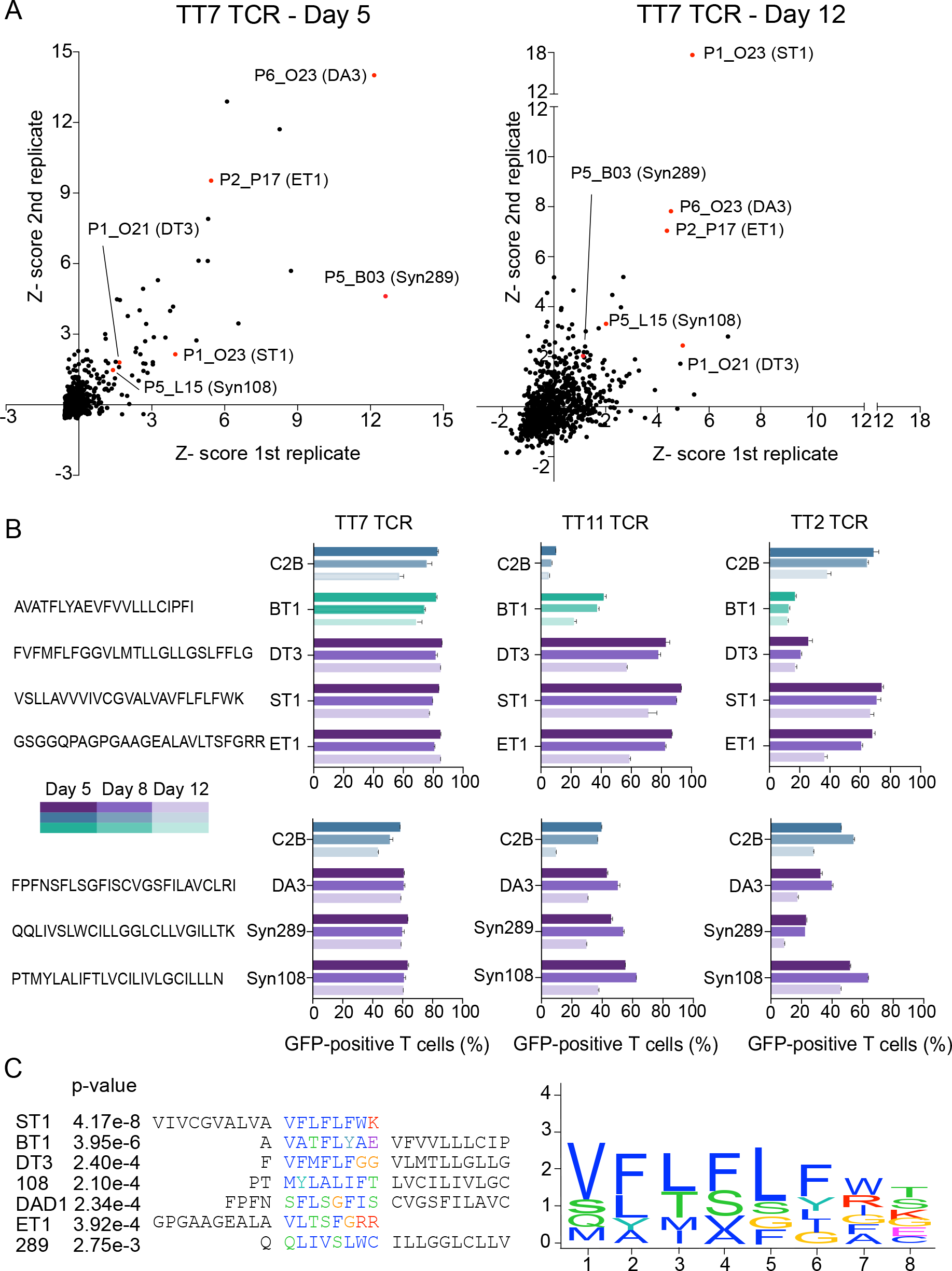
Functional identification of transmembrane domains for boosting class II presentation. (A) Results of a plate-based screen of ER_TM domain library elements using the TetX 916 – 1315 minigene and cognate TT7 TCR. Left panel, results obtained at day 5; right panel, results at day 12 post-transduction. The intensity of the GFP signal is represented as the Z-scores (over all tested constructs) of two replicates (x- and y-axes). Red dots indicate colonies selected for validation/follow-up. (B) Flow cytometric analysis of co-cultures (with their respective cognate TCRs) at day 5, day 8, and day 12 following transduction with the TetX 916 - 1315, TetX 612 - 1011, and TetX 1-399 minigenes. (C) Validated sequences from panel (B) were analyzed for a common motif using MEME-sorted from the MEME suite motif analysis tool set. The algorithm identified the ‘VFLFLFWK’ motif present in ST1 and partially conserved in the other validated sequences.

### PES fusion constructs can be used to boost and multiplex antigen processing

To ascertain the potential of the ST1 sequence as an antigen processing booster, we subcloned several minigenes containing the putative targets of class II restricted TCRs we had not yet been able to (or could only poorly) functionally validate with the T-FINDER platform, including with the BT1 fusion. We selected two COVID specific TCRs (4.1 and 6.3 ^32^) and minigenes containing a 400 amino acid fragment of the spike (S) protein of SARS-CoV-2 or its B1.525, B.1.1.529, and B.1.617.2 variants (Figure 4A), and observed a striking increase in T-cell activation upon N-terminal ST1 fusion. We also tested the ST1 construct with TCRs RA38 and RA48 from rheumatoid arthritis patients^33^, paired with their cognate minigene from the HCMV pp65 protein (Figure 4B). In both cases, ST1 fusion also increases the reporter response, from a weak signal barely indistinguishable from the negative control to a strong signal similar to that observed for the pulsed peptide control. Finally, we used the LS2.8/3.15, D2, and S16TCRs from celiac disease (CeD) patients^34, 35^ The TCRs specifically recognize a peptide from wheat gliadin (GDA9 protein), as well as a pathogen-derived mimicry epitope (g8pw65 protein). The CeD TCR reporter signals were low and difficult to discriminate from the negative control (GDA4 protein) when using unmodified or BT1-fused minigenes; however, when using the ST1 construct, T-cell activation was stronger to clearly identify a positive cognate interaction in T-FINDER (Figure 4C). In summary, the ST1 fusion construct improves the ability of our epitope identification platform to functionally validate or deconvolute TCRs of unknown specificity by increasing, both quantitatively and qualitatively, presented peptides derived from ectopically expressed putative antigen ORFs.

**Figure 4.**
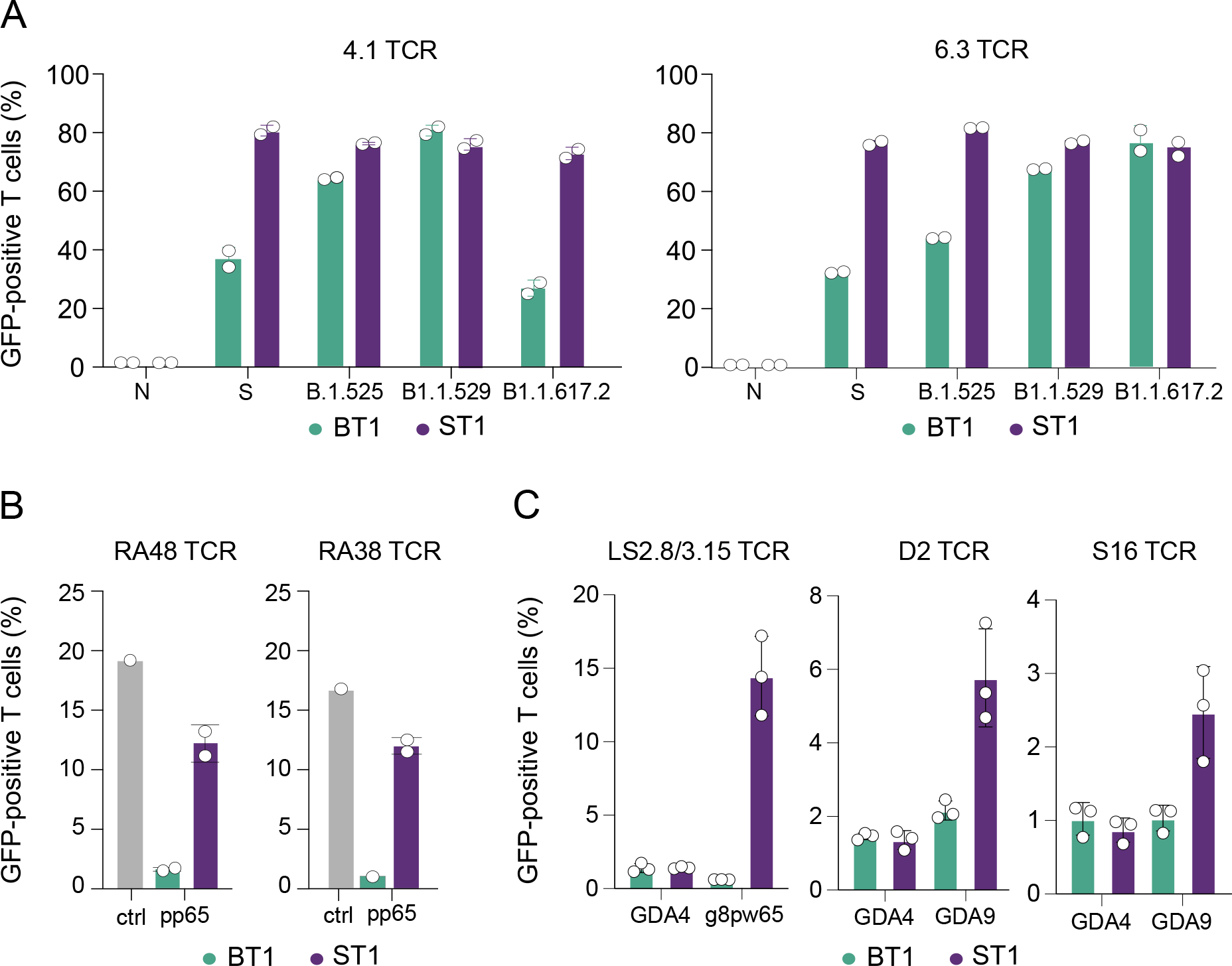
Validation of ST1 as an antigen presentation enhancing sequence. (A) Flow cytometric analysis of PNJE3 reporter cells expressing the TCRs 4.1 or 6.3 co-cultured with HLA-matched B-LCLs expressing spike protein from SARS-CoV-2 or B1.525, B.1.529, and B.1.617.2 variants cloned in the BT1 (green) or ST1 (violet) fusion constructs. Results are normalized to the negative control expressing a SARS-CoV-2 nucleocapsid (N) fragment. Percentage of GFP-positive reporter cells transduced with the RA38 or RA48 TCRs co-cultured with pulsed peptide (positive control) or the cognate pp65 1 - 399 minigene in the BT1 (green) or ST1 (violet) fusion constructs. (C) Reporter T-cell co-culture with celiac disease-derived TCRs (LS2.8/3.15, D2, and S16) with minigenes containing their respective cognate epitopes, GDA9 1 – 399 and G8PW65 1 - 399. The GDA4 1 – 399 minigene was used as a negative control.

## Discussion

The ability of EBV-immortalized B cells to process and present antigens encoded by a lentiviral vector on both class I and class II HLA complexes makes them an attractive option for antigen presenting cells in *in vitro* cellular assays. However, as demonstrated in Figure 1E, class II antigen processing capacity of transduced minigenes is rapidly lost over time. Previous descriptions of similar phenomena, such as the evasion of immunosurveillance by cancer through the downregulation of antigen presentation^36^, involve mechanisms such as genetic mutation, epigenetic silencing, or post transcriptional downregulation of the HLA loci^37^. In this work, we observed the attenuation of antigen presentation as an effect of impaired antigen processing, without perturbing the HLA presentation machinery, confirmed by the as pulsed peptide control. We observed that APA is stronger for class II antigens, suggesting a specific role for this process in regulating antigen presentation. Indeed, APA may reflect an additional mechanistic layer of enforcing T-cell peripheral tolerance. B cells are primarily known for their role in antibody production, and work synergistically with T cells to orchestrate adaptive immune responses^38^. B cells are also professional antigen presenting cells (they express class II HLA and can internalize cognate antigens via the BCR^39^) and are involved in the differentiation of CD4+ T cells. Based on these characteristics, APA may represent a potential role for B-cell involvement in controlling T-cell responses. In this model, once B cells have presented a “new” antigen to CD4+ T cells, their interaction with activated T cells leads to the downregulation of antigen processing, limiting the scope of their interaction to save resources and prevent potential autoimmune reactions. Further analyses in primary lymphocyte interactions are required to determine how this process is directed, whether it is antigen specific, and to identify the molecular pathways involved.

By functionally screening for epitope presentation (Figure 3), we identified several transmembrane domain sequences which overcame APA in our experimental setting; together with the TM1 of Bap31 (BT1), we identified 4 additional TM domains from ER proteins: the TM1 of SYT6, TM1 of ESYT1, TM3 of DERL3, and TM3 of DAD1, and 2 synthetic transmembrane domains. SYT6 is known to be involved in vesicular trafficking and exocytosis at synapses^40^, and ESYT1 is known to be involved in lipid transport^41^. DAD1^42^ functions as a complex subunit involved in protein glycosylation, while DERL3 was reported to regulate the ER-associated degradation (ERAD) of misfolded glycoproteins^43^. From this information it is difficult to identify a single functional pathway behind the role of the identified PES in antigen presentation, although a specific motif analysis of the positive PES comparing the 6 “hits” to the non-functional library sequences identified a core motif present in all 6 sequences (Figure 3C). Additional experiments are required to determine whether this motif is potentially a consensus binding region for another membrane-embedded protein or otherwise involved in protein trafficking or regulation of class II antigen processing, and to understand its mechanism for promoting strong and sustained presentation of epitopes from fused ORFs.

The results obtained in this work identify and overcome a crucial bottleneck in minigene-based antigen presentation platforms. Indeed, the antigen library presenting system developed here ensures strong and stable class I and class II antigen presentation from genetically encoded library elements. Furthermore, the ectopically expressed minigenes are naturally processed by the antigen-presenting cell, incorporating potential post-translational modifications^44, 45^ and *cis*-spliced peptides^10, 11^. The system is designed to be HLA independent (by using autologous or otherwise HLA-matched APC lines) and enables the screening of libraries covering more than 10^8^ bp of protein-coding genetic material across only 40,000 minigene elements. Coupled with our highly sensitive T-cell reporter^9^, these elements will greatly improve the success rate of unbiased antigen discovery, increasing our ability to explore the role of the adaptive immune system in autoimmune disease, cancer, and infectious disease.

## Acknowledgements

The authors would like to thank Nina Papavasiliou for her input and guidance throughout the project together with Nathan Felix and Murray McKinnon, and África Temporal Plo for her review of codon usage literature. This work was supported by BioMed X funding to JML from the Janssen Pharmaceutical Companies of Johnson and Johnson.

## Materials and Methods

### Lentiviral backbone construct and minigene cloning

The lentiviral backbone construct was designed starting from the commercial pLVX-EF1α-IRES-Puro vector (Takara Bio, 631253), to which a T2A-mTagBFP2 cassette (Twist Bioscience) was added. Each minigene was ordered as a gene fragment from Twist Bioscience, containing BamHI and SpeI restriction sites (Thermo Fisher Scientific GmbH, FD0055 and FD1254). Gene fragments were resuspended in TE Buffer (10 mM Tris-HCl, 0.1 mM EDTA), and 400 ng were digested (20 ul total volume). For PCR-amplified (e.g., subfragments of pre-existing constructs), gene-specific primers containing the restriction enzyme sequences were ordered from Intergate DNA technology (IDT) and resuspended at 100 µM with TE buffer. PCR reactions were performed using Phusion 5X Master mix (New England Biolabs GmbH, M0492S). For a 25 µl reaction, 12.5 µl of the master mix were used together with 500 nM of each primer and 1 ng of template. 10 µl of PCR reaction were digested in a 30 μl reaction with 1.2 µl of each enzyme and 2 µl Fast Digest buffer. Backbone preparation was performed in a 30 µl reaction with 3 µg of plasmid, 1.5 µl of each enzyme and 1 µl of alkaline phosphatase (Thermo Fisher Scientific GmbH MEF0651). Digests were incubated for 1h at 37 °C and then run on a 1% agarose gel for 15 min at 100 V. Gel extraction was performed following manufacturer protocols (Macherey-Nagel GmbH & Co. KG740609). Backbone ligation with the minigene was performed using 40 ng of backbone and a 3:1 molar ratio of the insert (Thermo Fisher Scientific GmbH EL0011). The ligation reaction was incubated for 2h at room temperature or overnight at 16°C. 18 µl of Stellar (Takara Bio Europe SAS 636766) chemically competent bacteria were transformed using 2 µl of the ligation reaction. After initial incubation on ice for at least 45 min, a heat shock was performed at 42 °C for 45 sec, followed by a short incubation on ice for 2-3 minutes. Bacteria were diluted with 100 µl of SOC and incubated at 37 °C for 1h, then plated on LB agar plates containing 100 µg/ml ampicillin; plates were incubated for 20h to 24h at 32 °C before sending isolated colonies for Sanger sequencing. Colonies were sequenced using E. coli NightSeq sequencing service (Microsynth AG); positive colonies were cultured in 100 ml of 2YT medium and prepped using a midi-scale commercial kit (Macherey-Nagel 740410100).

### Codon-optimized backtranslation

The FALCON script is available at https://github.com/lindnerIAA/FALCON.

### Cell culture

HEK293FT cells were cultured in DMEM (Thermo Fisher Scientific GmbH 61965059) with 10% FBS and 1% penicillin-streptomycin (Capricorn Scientific PS-B), at 37°C in a humidified 5% atmosphere. BOLETH cells were cultured in RMPI 1640 Medium (Thermo Fisher Scientific GmbH 61870044) supplemented with 10% FBS (PAN-Biotech GmbHP30-3306), 1% penicillin-streptomycin (Capricorn Scientific PS-B), 50 mM β-mercaptoethanol (Fisher Scientific GmbH 21985023), 1 mM sodium pyruvate (Thermo Fisher Scientific GmbH 11360039), and 5ml MEM Non-Essential Amino Acids (Thermo Fisher Scientific GmbH 11140035) at 37°C in a humidified 5% CO_2_ atmosphere. TCR-deficient Jurkat cells were cultured in RPMI 1640 Medium + 10% heat-inactivated (56°C for 20 min) FBS and 1% penicillin-streptomycin at 37 °C in a humidified 5% CO2 atmosphere. Jurkat cells expressing the TCR reporter construct were maintained in culture with 10 µg/ml blasticidin (Invivogen ant-bl-1). For TCR expression, cells were selected with 1 µg/ml puromycin (Invivogen ant-pr-1).

### Lentiviral production and transduction

HEK293T cells were used for lentiviral production; the day before transfection 3×10^5^ cells were plated in 6-well plates in 2 ml of medium. For each transfection, 0.67 µg of PMD2.G 1µg of PAX2 lentiviral packaging plasmids together with 1.3 µg of minigene plasmid were used. DNA was mixed in 200 µl of Optimem (Gibco 31985062) before adding 6 µl of TransIT VirusGen transfection reagent (Mirus MIR6700). The transfection mixture was incubated for 10 to 15 min and then added dropwise to HEK293T cells. 48h after transfection, viral supernatant was collected and filtered with a 0.45 µm syringe filter (Th. Geyer GmbH & Co. KG 16537K). BOLETH cells were treated before transduction with 2 µM BX-795-hydrochloride (Sigma-Aldrich Chemie GmbH SML0694-25MG) for 4h. 1.5×10^6^ cells were washed and resuspended in 1 ml of BOLETH medium, the filtered virus was added dropwise together with 7μg/ml polybrene and 2 µM BX-795-hydrochloride. For T cells, only polybrene (Thermo Fisher Scientific GmbH 11360039) was added to the transduction. After 24h, cells were washed, and medium was replaced without additives. BOLETH cells were used for the experiments at day 5 post-transduction. T-cell reporter cell lines transduced with TCR were placed in selection medium 5 days post-transduction.

### B:T-cell co-cultures

BOLETH and T cells used in the co-culture experiments were counted and resuspended to a final concentration of 10^5^ cells/40 µl for BOLETH cells and 10^5^ cells/35 µl for T cells in BOLETH medium. Cells were plated in 96-well round bottom plates (neolab Migge GmbH C-8207). For positive controls, peptides were added to a final concentration of 1µM for tetanospasmin-derived peptides or 10 µM for class I peptides. In the case of unknown peptides, positive control wells were pre-treated with anti-CD3 for 2h prior to the experiment. The experiment was incubated for 16h at 37°C in a humidified 5% CO_2_ atmosphere before FACS staining. For longitudinal experiments, the co-culture was plated at day 5, day 8, and day 12 after transduction using the same batch of transduced cells.

### Microscope-based screening

The TM_ER library was screened, recording the sfGFP signal on a Nikon Eclipse Ti2 microscope. For the experiment, the HEK293 transfection and BOLETH transduction were performed in a 96-well plate, while the experiment was plated in a 384-well plate (Cell culture microplate 384 well black, Greiner Bio-one GmbH). Colonies were picked in a deep 96-well reservoir (with 1.5 ml of 2xYT medium plus relevant antibiotics). The plate was covered with a breathable seal and incubated for 20 h at 32 °C with mild shaking (90 rpm). DNA minipreps were performed following manufacturer guidelines according to the centrifuge protocol (NZYTech MB20201). Both transduction and transfection protocols follow the guidelines described before, but with some adjustments for the well format. For each transfection, 3.5×10^4^ HEK cells were plated in 96-well, flat-bottom plates in 210 µl of DMEM. 24h later, they were transfected with 10 µl of Optimem 50 ng of PAX, 33.5 ng of pMDG, 83 ng of plasmid and 0.3 µl of TransIT. To facilitate the pipetting, first the DNA was added together with 5 µl of Optimem and the remaining 5 µl were added together with the TransIT. The transduction was performed 48h after the transfection, BOLETH cells were pre-treated with BX-795-HCl and for each transduction, 5×105 BOLETH cells in 60 µl of medium were used. HEK293 plates were centrifuged 5 min at 800 rpm, and 60 to 70 µl of viral supernatant were added to the BOLETH cells together with 2 µM BX-795-hydrochloride and 7μg/ml polybrene. 24h after transduction, the BOLETH medium was replaced with 140 µl of fresh medium. First, the medium was replaced with 100 µl per well of DMEM Fluorobrite (Thermo Fisher Scientific GmbHA1896701) (5% FBS, β-mercaptoethanol, NEAA, sodium pyruvate, and penicillin/streptomycin), and BOLETH clumps were disassociated mechanically. 40 µl of BOLETH cells were moved to a 384-well plate (Greiner Bio-one GmbH 781091), the experiment was plated in technical duplicates. 2.5×10^5^ T cells in 35 µl of DMEM Fluorobrite were then added to each well. After 7h, the sfGFP signal was measured using exposure 600 ms and 10X magnification. Image analysis was performed with ImageJ and the results were plotted as transformed *Z* scores.

### FACS sample staining analysis

Samples were diluted in FACS buffer (PBS with 2mM EDTA and 2% FBS) and washed twice before staining with 50 µl of FACS buffer containing antibodies diluted 1:200 and incubated at 4 °C for 20 minutes. Samples were washed twice and resuspended in a final volume of 100 µl of FACS buffer. Sample measurements were recorded using a FACS Aria II (BD Biosciences), and data were analyzed with FlowJo. In the analysis, the percentage of positive cells in the negative control was always set to 1%.

### Antigen processing signal fusion sequences

CMA

MKETAAAKFERQHMDSS

ATGAAAGAGACCGCCGCAGCCAAGTTCGAAAGACAGCACATGGACTCTAGCA

DC-LAMP

N-term

MPRQLRAAAALFARLAVILH

ATGCCGAGGCAGTTGAGGGCGGCGGCGGCGTTGTTTGCGAGGTTGGCGGTGATATT GCAT

C-term

SSDYTIVLPVIGAIVVGLCLMGMGVYKIRLRCQSSGYQRI

AGCAGCGATTATACGATAGTGTTGCCGGTGATAGGGGCGATAGTGGTGGGGTTGTGTTTGATGGGGATGGGGGTGTATAAGATAAGGTTGAGGTGTCAGAGCAGCGGGTATCAGAGGATA

LIR-ATG14

TDLGTDWENLPSPRFC

ACCGACTTAGGCACCGACTGGGAAAATTTACCCTCCCCCAGATTTTGT

Bcap31_TM1_21_7 (BT1)

AVATFLYAEVFVVLLLCIPFI

GCAGTTGCCACCTTCCTCTATGCGGAGGTCTTTGTTGTGTTGCTTCTCTGCATTCCCT TCATT

ESYT1_37_25 (ET1)

GSGGQPAGPGAAGEALAVLTSFGRR

GGTTCTGGCGGGCAACCAGCAGGACCAGGAGCAGCTGGAGAAGCCCTGGCTGTTCTGACTTCTTTTGGAAGAAGA

SYT6_58_25 (ST1)

VSLLAVVVIVCGVALVAVFLFLFWK

GTTTCCTTGCTGGCCGTTGTAGTAATCGTGTGTGGGGTGGCTCTCGTAGCAGTTTTCCTGTTCCTGTTCTGGAAG

DERL3_97_25 (DT3)

FVFMFLFGGVLMTLLGLLGSLFFLG

TTCGTTTTCATGTTCCTGTTCGGGGGCGTTCTGATGACTCTGCTGGGGTTGCTGGGAAGCCTGTTCTTCCTGGGA

>DAD1_51_25 (DT1)

FPFNSFLSGFISCVGSFILAVCLRI

TTCCCTTTCAATTCCTTCCTGAGCGGCTTCATCTCGTGCGTCGGCAGCTTTATTCTGGCAGTGTGCCTGCGCATT

Synthetic_298

QQLIVSLWCILLGGLCLLVGILLTK

CAACAGCTGATTGTCTCTCTCTGGTGCATTCTGCTGGGCGGCCTCTGCCTGCTGGTGGGAATTCTGCTGACTAAG

Synthetic_108

PTMYLALIFTLVCILIVLGCILLLN

CCAACAATGTACCTGGCCCTGATCTTTACCCTGGTGTGTATCCTGATCGTGCTGGGATGCATCCTGCTGCTGAAC

CD74

N-term

MHRRRSRSCREDQKPVMDDQRDLISNNEQLPMLGRRPGAPESKCSRGALYTGFSILVTLLLAGQATTAYFLYQQQGRLDKLTVTSQNLQLENLRMKLPKPPKPVSK

ATGCACCGTAGGCGGTCAAGATCATGCAGGGAAGATCAGAAGCCGGTAATGGACGACCAGCGGGACCTGATTAGCAACAATGAGCAGCTGCCCATGCTCGGCAGACGACCCGGGGCTCCAGAGAGTAAATGTTCTAGGGGCGCTCTTTACACCGGGTTTAGTATTCTTGTGACTTTATTACTGGCTGGGCAGGCTACTACCGCATATTTCCTGTACCAACAGCAGGGCCGTCTGGATAAGCTGACAGTGACATCACAGAACTTGCAGCTCGAGAACCTGAGAATGAAGCTCCCGAAGCCTCCGAAGCCTGTTTCGAAA

C-term

QALPMGALPQGPMQNATKYGNMTEDHVMHLLQNADPLKVYPPLKGSFPENLRHLKNTM ETIDWKVFESWMHHWLLFEMSRHSLEQKPTDAPPKESLELEDPSSGLGVTKQDLGPVP M

CAGGCCCTGCCCATGGGAGCCCTCCCCCAGGGCCCCATGCAGAACGCCACCAAGTACGGCAACATGACCGAGGACCATGTGATGCATCTGCTGCAGAACGCCGATCCTCTGAAGGTGTACCCACCGCTGAAGGGGTCCTTCCCAGAGAACCTTAGACATTTGAAGAACACGATGGAGACCATCGATTGGAAGGTCTTTGAATCCTGGATGCATCACTGGCTGCTGTTCGAGATGTCCAGGCACTCCCTTGAGCAGAAGCCCACAGACGCCCCGCCTAAAGAGTCCTTGGAGTTAGAGGACCCATCCTCTGGTCTTGGAGTCACGAAGCAGGATCTCGGACCTGTGCCAATG

### List of minigenes

MLANA A27L

MPREDAHFIYGYPKKGHGHSYTTAEELAGIGILTVILGVLLLIGCWYCRRRNGYRALMDKSLHVGTQCALTRRCPQEGFDHRDSKVSLQEKNCEPVVPNAPPAYEKLSAEQSPPPYSP

ATGCCCAGGGAGGACGCCCACTTCATCTACGGCTACCCCAAGAAGGGCCACGGCCACAGCTACACCACTGCAGAGGAGCTTGCTGGGATCGGCATCCTGACAGTGATCCTAGGCGTGCTGCTGCTGATCGGCTGCTGGTACTGCCGGAGGAGGAACGGCTACAGGGCCCTGATGGACAAGAGCCTGCACGTGGGCACCCAGTGCGCCCTGACCAGGAGGTGCCCCCAGGAGGGCTTCGACCACAGGGACAGCAAGGTGAGCCTCCAGGAGAAGAACTGCGAGCCCGTGGTGCCCAACGCCCCCCCCGCCTACGAGAAGCTGAGCGCCGAGCAGAGCCCTCCACCATACAGCCCC

TetX 1 - 399

MPITINNFRYSDPVNNDTIIMMEPPYCKGLDIYYKAFKITDRIWIVPERYEFGTKPEDFNPPSSLIEGASEYYDPNYLRTDSDKDRFLQTMVKLFNRIKNNVAGEALLDKIINAIPYLGNSYSLLDKFDTNSNSVSFNLLEQDPSGATTKSAMLTNLIIFGPGPVLNKNEVRGIVLRVDNKNYFPCRDGFGSIMQMAFCPEYVPTFDNVIENITSLTIGKSKYFQDPALLLMHELIHVLHGLYGMQVSSHEIIPSKQEIYMQHTYPISAEELFTFGGQDANLISIDIKNDLYEKTLNDYKAIANKLSQVTSCNDPNIDIDSYKQIYQQKYQFDKDSNGQYIVNEDKFQILYNSIMYGFTEIELGKKFNIKTRLSYFSMNHDPVKIPNLLDDTIYNDTEGF

ATGCCCATCACAATCAATAACTTTCGGTACTCAGACCCTGTAAATAACGACACTATTATCATGATGGAGCCACCCTACTGCAAGGGGTTGGACATCTATTACAAGGCCTTTAAGATCACAGACAGGATTTGGATCGTGCCAGAGCGATACGAGTTCGGCACCAAGCCCGAAG

ACTTCAACCCTCCCAGCAGCCTGATTGAGGGTGCCTCCGAGTACTACGACCCTAATTACCTGAGAACCGATTCCGACAAGGATAGATTCCTCCAGACCATGGTCAAGCTGTTCAATAGGATAAAGAATAATGTCGCAGGTGAGGCCCTGCTGGATAAGATCATTAACGCGATCCCTTATCTGGGGAACTCTTACAGTTTGCTCGACAAGTTCGACACAAACAGCAACTCGGTATCCTTCAACCTTCTGGAACAGGACCCAAGCGGCGCCACGACTAAGTCGGCCATGCTCACAAATCTCATTATTTTCGGGCCAGGGCCCGTCTTGAATAAGAACGAAGTAAGGGGGATCGTCCTGCGGGTGGATAACAAGAACTACTTTCCGTGCCGGGATGGATTTGGGAGCATTATGCAGATGGCTTTTTGTCCGGAGTATGTGCCCACGTTCGACAATGTCATCGAGAACATAACCTCCCTGACCATTGGTAAATCTAAGTACTTCCAAGACCCCGCTCTGCTTCTGATGCATGAGCTCATCCATGTTCTCCATGGCTTGTATGGAATGCAGGTATCCTCGCACGAAATTATTCCCAGTAAGCAGGAGATCTATATGCAGCACACATACCCCATCTCAGCTGAAGAGCTGTTCACGTTCGGGGGTCAGGACGCCAATCTGATCTCCATCGACATTAAGAACGACCTATATGAAAAGACCCTAAACGATTATAAGGCCATTGCGAACAAGCTGAGCCAGGTCACCAGCTGTAATGACCCTAATATCGACATTGACTCTTACAAGCAGATTTATCAGCAGAAGTACCAGTTCGATAAGGATAGCAACGGGCAGTACATTGTCAACGAGGACAAATTTCAGATCCTCTACAATTCTATCATGTACGGGTTCACAGAAATCGAGCTCGGCAAAAAGTTCAACATAAAGACACGTCTGTCCTACTTCTCCATGAATCACGATCCCGTGAAGATCCCTAATCTGCTGGACGACACCATCTATAACGACACCGAAGGCTTC

TetX 50 - 149

TACGAATTTGGCACAAAGCCAGAGGACTTTAATCCGCCCAGTTCCCTGATAGAGGGCGCCTCCGAATATTACGACCCCAACTATCTGCGGACTGACTCCGACAAGGACCGCTTTCTGCAGACTATGGTTAAGTTATTCAACCGCATTAAAAACAATGTCGCCGGTGAGGCGTTGCTCGACAAAATCATCAACGCCATCCCCTACTTGGGCAACTCTTACTCGCTGCTTGATAAATTCGACACCAACAGCAATAGTGTTAGCTTTAACCTGCTGGAGCAGGACCCGTCGGGCGCTACTACC YEFGTKPEDFNPPSSLIEGASEYYDPNYLRTDSDKDRFLQTMVKLFNRIKNNVAGEALLDKIINAIPYLGNSYSLLDKFDTNSNSVSFNLLEQDPSGATT

TetX 31 - 155

GCATTCAAAATCACTGACCGGATATGGATTGTGCCCGAGAGATACGAATTTGGCACAAAGCCAGAGGACTTTAATCCGCCCAGTTCCCTGATAGAGGGCGCCTCCGAATATTACGACCCCAACTATCTGCGGACTGACTCCGACAAGGACCGCTTTCTGCAGACTATGGTTAAGTTATTCAACCGCATTAAAAACAATGTCGCCGGTGAGGCGTTGCTCGACAAAATCATCAACGCCATCCCCTACTTGGGCAACTCTTACTCGCTGCTTGATAAATTCGACACCAACAGCAATAGTGTTAGCTTTAACCTGCTGGAGCAGGACCCGTCGGGCGCTACTACCAAGAGTGCCATGCTCACCAACCTCATAATTTTC

AFKITDRIWIVPERYEFGTKPEDFNPPSSLIEGASEYYDPNYLRTDSDKDRFLQTMVKLFNRIKNNVAGEALLDKIINAIPYLGNSYSLLDKFDTNSNSVSFNLLEQDPSGATTKSAMLTNLII F

TetX 22 - 171

TGTAAGGGGTTGGACATTTATTATAAAGCATTCAAAATCACTGACCGGATATGGATTGTGCCCGAGAGATACGAATTTGGCACAAAGCCAGAGGACTTTAATCCGCCCAGTTCCCTGATAGAGGGCGCCTCCGAATATTACGACCCCAACTATCTGCGGACTGACTCCGACAAGGACCGCTTTCTGCAGACTATGGTTAAGTTATTCAACCGCATTAAAAACAATGTCGCCGGTGAGGCGTTGCTCGACAAAATCATCAACGCCATCCCCTACTTGGGCAACTCTTACTCGCTGCTTGATAAATTCGACACCAACAGCAATAGTGTTAGCTTTAACCTGCTGGAGCAGGACCCGTCGGGCGCTACTACCAAGAGTGCCATGCTCACCAACCTCATAATTTTCGGGCCTGGGCCCGTTCTGAACAAAAATGAGGTCCGTGGTATTGTACTC

CKGLDIYYKAFKITDRIWIVPERYEFGTKPEDFNPPSSLIEGASEYYDPNYLRTDSDKDRFLQTMVKLFNRIKNNVAGEALLDKIINAIPYLGNSYSLLDKFDTNSNSVSFNLLEQDPSGATTKSAMLTNLIIFGPGPVLNKNEVRGIVL

TetX 15 - 179

ATTATGATGGAGCCCCCTTACTGTAAGGGGTTGGACATTTATTATAAAGCATTCAAAATCACTGACCGGATATGGATTGTGCCCGAGAGATACGAATTTGGCACAAAGCCAGAGGACTTTAATCCGCCCAGTTCCCTGATAGAGGGCGCCTCCGAATATTACGACCCCAACTATCTGCGGACTGACTCCGACAAGGACCGCTTTCTGCAGACTATGGTTAAGTTATTCAACCGCATTAAAAACAATGTCGCCGGTGAGGCGTTGCTCGACAAAATCATCAACGCCATCCCCTACTTGGGCAACTCTTACTCGCTGCTTGATAAATTCGACACCAACAGCAATAGTGTTAGCTTTAACCTGCTGGAGCAGGACCCGTCGGGCGCTACTACCAAGAGTGCCATGCTCACCAACCTCATAATTTTCGGGCCTGGGCCCGTTCTGAACAAAAATGAGGTCCGTGGTATTGTACTCAGAGTCGACAATAAAAACTACTTCCCCTGCCGGGACGGTTTCGGGAGCATAATG

IMMEPPYCKGLDIYYKAFKITDRIWIVPERYEFGTKPEDFNPPSSLIEGASEYYDPNYLRTDSDKDRFLQTMVKLFNRIKNNVAGEALLDKIINAIPYLGNSYSLLDKFDTNSNSVSFNLLEQDPSGATTKSAMLTNLIIFGPGPVLNKNEVRGIVLRVDNKNYFPCRDGFGSIM

TetX 1 - 199

AACAATTTCCGTTATAGCGATCCCGTGAACAACGACACGATTATTATGATGGAGCCCCCTTACTGTAAGGGGTTGGACATTTATTATAAAGCATTCAAAATCACTGACCGGATATGGATTGTGCCCGAGAGATACGAATTTGGCACAAAGCCAGAGGACTTTAATCCGCCCAGTTCCCTGATAGAGGGCGCCTCCGAATATTACGACCCCAACTATCTGCGGACTGACTCCGACAAGGACCGCTTTCTGCAGACTATGGTTAAGTTATTCAACCGCATTAAAAACAATGTCGCCGGTGAGGCGTTGCTCGACAAAATCATCAACGCCATCCCCTACTTGGGCAACTCTTACTCGCTGCTTGATAAATTCGACACCAACAGCAATAGTGTTAGCTTTAACCTGCTGGAGCAGGACCCGTCGGGCGCTACTACCAAGAGTGCCATGCTCACCAACCTCATAATTTTCGGGCCTGGGCCCGTTCTGAACAAAAATGAGGTCCGTGGTATTGTACTCAGAGT

CGACAATAAAAACTACTTCCCCTGCCGGGACGGTTTCGGGAGCATAATGCAGATGGCCTTCTGCCCAGAGTACGTG

NNFRYSDPVNNDTIIMMEPPYCKGLDIYYKAFKITDRIWIVPERYEFGTKPEDFNPPSSLIEGASEYYDPNYLRTDSDKDRFLQTMVKLFNRIKNNVAGEALLDKIINAIPYLGNSYSLLDKFDTNSNSVSFNLLEQDPSGATTKSAMLTNLIIFGPGPVLNKNEVRGIVLRVDNKNYFPCRDGFGSIMQMAFCPEYV

TetX 612 - 1011

LFLQWVRDIIDDFTNESSQKTTIDKISDVSTIVPYIGPALNIVKQGYEGNFIGALETTGVVLLLEYIPEITLPVIAALSIAESSTQKEKIIKTIDNFLEKRYEKWIEVYKLVKAKWLGTVNTQFQKRSYQMYRSLEYQVDAIKKIIDYEYKIYSGPDKEQIADEINNLKNKLEEKANKAMININIFMRESSRSFLVNQMINEAKKQLLEFDTQSKNILMQYIKANSKFIGITELKKLESKINKVFSTPIPFSYSKNLDCWVDNEEDIDVILKKSTILNLDINNDIISDISGFNSSVITYPDAQLVPGINGKAIHLVNNESSEVIVHKAMDIEYNDMFNNFTVSFWLRVPKVSASHLEQYGTNEYSIISSMKKHSLSIGSGWSVSLKGNNLIWTLKDSAGE

CTGTTCCTCCAGTGGGTCCGAGACATCATCGATGACTTCACGAACGAGTCTTCGCAGAAGACTACGATAGACAAGATCTCCGACGTGAGCACAATTGTTCCCTACATCGGGCCCGCTTTGAACATCGTGAAGCAGGGCTACGAGGGGAACTTTATCGGTGCACTTGAAACTACGGGTGTGGTGCTGCTCCTGGAGTATATCCCGGAGATCACCCTCCCAGTCATCGCCGCCCTGAGCATTGCCGAGAGCTCCACCCAAAAAGAGAAGATTATTAAAACCATAGACAACTTCCTGGAGAAGCGCTACGAGAAATGGATTGAGGTTTATAAGCTGGTAAAGGCCAAATGGCTAGGGACCGTCAACACCCAGTTTCAGAAGCGGAGCTACCAGATGTACCGCTCTCTGGAGTACCAGGTCGACGCGATTAAGAAGATCATTGATTACGAGTACAAAATCTACAGCGGACCGGACAAAGAGCAGATTGCGGACGAAATCAACAACCTAAAAAACAAGCTGGAGGAGAAGGCAAACAAGGCCATGATTAACATCAACATCTTCATGAGGGAATCTAGCCGGTCCTTCCTCGTGAATCAGATGATAAACGAAGCAAAAAAGCAGCTCCTGGAATTTGACACTCAGAGCAAGAACATACTGATGCAATACATTAAGGCTAACTCCAAGTTCATCGGGATCACTGAGCTGAAAAAACTTGAGTCTAAAATAAACAAGGTTTTTTCAACCCCAATACCATTCAGCTACTCCAAGAATCTGGACTGCTGGGTGGATAACGAAGAGGACATAGACGTGATCCTAAAGAAGAGTACAATCCTAAATCTGGACATAAACAATGACATCATCTCCGATATAAGCGGCTTCAACTCCTCCGTAATAACCTATCCTGACGCCCAACTGGTGCCCGGCATCAACGGTAAAGCCATACATCTGGTCAACAATGAGTCCAGCGAGGTGATCGTGCACAAGGCTATGGATATTGAGTATAACGACATGTTCAACAATTTCACCGTGTCCTTCTGGTTGAGGGTGCCAAAGGTCTCTGCTAGCCATCTGGAACAGTATGGGACCAACGAATACTCCATCATCAGTAGCATGAAGAAGCACAGCCTGTCCATTGGGTCCGGCTGGAGCGTGTCTCTGAAAGGCAACAACTTGATTTGGACCCTCAAGGACAGTGCCGGGGAA

TetX 916 - 1315

LFLQWVRDIIDDFTNESSQKTTIDKISDVSTIVPYIGPALNIVKQGYEGNFIGALETTGVVLLLEYIPEITLPVIAALSIAESSTQKEKIIKTIDNFLEKRYEKWIEVYKLVKAKWLGTVNTQFQKRS

YQMYRSLEYQVDAIKKIIDYEYKIYSGPDKEQIADEINNLKNKLEEKANKAMININIFMRESSRSFLVNQMINEAKKQLLEFDTQSKNILMQYIKANSKFIGITELKKLESKINKVFSTPIPFSYSKNLDCWVDNEEDIDVILKKSTILNLDINNDIISDISGFNSSVITYPDAQLVPGINGKAIHLVNNESSEVIVHKAMDIEYNDMFNNFTVSFWLRVPKVSASHLEQYGTNEYSIISSMKKHSLSIGSGWSVSLKGNNLIWTLKDSAGE

CCCGGGATCAATGGTAAGGCTATACATCTCGTCAATAACGAGTCCTCTGAGGTGATTGTGCATAAGGCAATGGACATTGAGTACAACGATATGTTCAACAATTTCACAGTGAGTTTCTGGCTGAGGGTCCCAAAAGTCTCGGCCTCCCACCTGGAGCAATACGGGACCAATGAGTATTCTATAATTAGCTCTATGAAGAAACACTCGTTGAGCATCGGAAGTGGCTGGAGTGTCAGCCTCAAGGGCAACAACCTCATCTGGACTCTGAAGGATTCTGCAGGGGAAGTGCGGCAGATTACTTTCAGGGACCTGCCCGATAAATTCAACGCCTACTTGGCTAATAAGTGGGTCTTTATTACGATCACCAACGATCGGTTGAGCTCTGCCAATCTCTACATCAACGGGGTGTTGATGGGCTCCGCTGAAATAACCGGCCTAGGCGCCATCCGGGAAGACAACAACATTACCCTCAAGTTAGACCGATGCAACAACAACAACCAGTATGTTAGCATCGATAAATTTCGGATCTTCTGTAAGGCGCTTAATCCGAAGGAAATCGAAAAACTTTACACCAGCTATCTTAGCATCACATTCCTGAGGGACTTCTGGGGGAACCCCCTTCGCTATGACACCGAGTATTATCTAATTCCCGTTGCTAGCTCTTCTAAGGACGTCCAGCTGAAGAACATCACGGACTATATGTACTTGACGAATGCGCCGTCGTACACCAACGGCAAGCTCAACATTTACTATCGACGGCTGTATAACGGCCTGAAGTTCATTATAAAGAGGTACACTCCCAATAATGAGATCGATTCTTTCGTTAAAAGCGGTGATTTCATCAAGCTGTACGTCAGCTATAACAATAACGAGCATATCGTCGGCTACCCGAAAGACGGAAACGCCTTCAATAACCTGGACCGCATCTTGCGTGTTGGGTATAATGCCCCGGGCATCCCCCTTTATAAGAAGATGGAGGCGGTGAAGTTGCGCGACCTCAAGACCTATTCGGTCCAGCTGAAATTGTACGATGACAAGAACGCATCTCTGGGGCTCGTCGGAACACACAACGGGCAAATAGGCAATGACCCCAATCGGGACATCCTCATAGCTTCGAATTGGTACTTTAACCATCTGAAGGACAAAATTCTGGGATGCGACTGGTACTTCGTGCCGACTGACGAGGGCTGGACTAACGAC

N_400_2

SDNGPQNQRNAPRITFGGPSDSTGSNQNGERSGARSKQRRPQGLPNNTASWFTALTQHGKEDLKFPRGQGVPINTNSSPDDQIGYYRRATRRIRGGDGKMKDLSPRWYFYYLGTGPEAGLPYGANKDGIIWVATEGALNTPKDHIGTRNPANNAAIVLQLPQGTTLPKGFYAEGSRGGSQASSRSSSRSRNSSRNSTPGSSRGTSPARMAGNGGDAALALLLLDRLNQLESKMSGKGQQQQGQTVTKKSAAEASKKPRQKRTATKAYNVTQAFGRRGPEQTQGNFGDQELIRQGTDYKHWPQIAQFAPSASAFFGMSRIGMEVTPSGTWLTYTGAIKLDDKDPNFKDQVILLNKHIDAYKTFPPTEPKKDKKKKADETQALPQRQKKQQTVTLLPAADLD

S_400_100

IIRGWIFGTTLDSKTQSLLIVNNATNVVIKVCEFQFCNDPFLGVYYHKNNKSWMESEFRVYSSANNCTFEYVSQPFLMDLEGKQGNFKNLREFVFKNIDGYFKIYSKHTPINLVRDLPQGFS

ALEPLVDLPIGINITRFQTLLALHRSYLTPGDSSSGWTAGAAAYYVGYLQPRTFLLKYNENGTITDAVDCALDPLSETKCTLKSFTVEKGIYQTSNFRVQPTESIVRFPNITNLCPFGEVFNATRFASVYAWNRKRISNCVADYSVLYNSASFSTFKCYGVSPTKLNDLCFTNVYADSFVIRGDEVRQIAPGQTGKIADYNYKLPDDFTGCVIAWNSNNLDSKVGGNYNYLYRLFRKSNLKPFERDISTEIYQAGSTPCNGVEGFNCYFPLQSYGFQP

S_B.1.617.2_400_100

IIRGWIFGTTLDSKTQSLLIVNNATNVVIKVCEFQFCNDPFLDVYYHKNNKSWMESGVYSSANNCTFEYVSQPFLMDLEGKQGNFKNLREFVFKNIDGYFKIYSKHTPINLVRDLPQGFSALEPLVDLPIGINITRFQTLLALHRSYLTPGDSSSGWTAGAAAYYVGYLQPRTFLLKYNENGTITDAVDCALDPLSETKCTLKSFTVEKGIYQTSNFRVQPTESIVRFPNITNLCPFGEVFNATRFASVYAWNRKRISNCVADYSVLYNSASFSTFKCYGVSPTKLNDLCFTNVYADSFVIRGDEVRQIAPGQTGKIADYNYKLPDDFTGCVIAWNSNNLDSKVGGNYNYRYRLFRKSNLKPFERDISTEIYQAGSKPCNGVEGFNCYFPLQSYGFQPTN

S_B.1.1.529_400_100

RGWIFGTTLDSKTQSLLIVNNATNVVIKVCEFQFCNDPFLDHKNNKSWMESEFRVYSSANNCTFEYVSQPFLMDLEGKQGNFKNLREFVFKNIDGYFKIYSKHTPIIVEPERDLPQGFSALEPLVDLPIGINITRFQTLLALHRSYLTPGDSSSGWTAGAAAYYVGYLQPRTFLLKYNENGTITDAVDCALDPLSETKCTLKSFTVEKGIYQTSNFRVQPTESIVRFPNITNLCPFDEVFNATRFASVYAWNRKRISNCVADYSVLYNLAPFSTFKCYGVSPTKLNDLCFTNVYADSFVIRGDEVRQIAPGQTGNIADYNYKLPDDFTGCVIAWNSNKLDSKVSGNYNYLYRLFRKSNLKPFERDISTEIYQAGNKPCNGVAGFNCYFPLRSYSFRPTYG

S_B1.525_400_100

RGWIFGTTLDSKTQSLLIVNNATNVVIKVCEFQFCNDPFLGVYHKNNKSWMESEFRVYSSANNCTFEYVSQPFLMDLEGKQGNFKNLREFVFKNIDGYFKIYSKHTPINLVRDLPQGFSALEPLVDLPIGINITRFQTLLALHRSYLTPGDSSSGWTAGAAAYYVGYLQPRTFLLKYNENGTITDAVDCALDPLSETKCTLKSFTVEKGIYQTSNFRVQPTESIVRFPNITNLCPFGEVFNATRFASVYAWNRKRISNCVADYSVLYNSASFSTFKCYGVSPTKLNDLCFTNVYADSFVIRGDEVRQIAPGQTGKIADYNYKLPDDFTGCVIAWNSNNLDSKVGGNYNYLYRLFRKSNLKPFERDISTEIYQAGSTPCNGVKGFNCYFPLQSYGFQPTNG pp65_308_253

MTMTRNPQPFMRPHERNGFTVLCPKNMIIKPGKISHIMLDVAFTSHEHFGLLCPKSIPGLSISGNLLMNGQQIFLEVQAIRETVELRQYDPVAALFFFDIDLLLQRGPQYSEHPTFTSQYRIQGKLEYRHTWDRHDEGAAQGDDDVWTSGSDSDEELVTTERKTPRVTGGGAMAGASTSAGRKRKSASSATACTSGVMTRGRLKAESTVAPEEDTDEDSDNEIHNPAVFTWPPWQAGILARNLVPMVATVQGQNLKYQEFFWDANDIYRIFAELEGVWQPAAQPKRRRHRQDALPGPCIASTPKKHRG

GDA4_400_2

KTFLILALRAIVATTATIAVRVPVPQLQPQNPSQQQPQKQVPLVQQQQFPGQQQPFPPQQPYPQQQPFPSQQPYMQLQPFPQPQLPYPQPQLPYPQPQPFRPQQSYPQPQPQYSQPQQPISQQQQQQQQQQQQQQQILQQILQQQLIPCRDVVLQQHSIAHGSSQVLQQSTYQLVQQFCCQQLWQIPEQSRCQAIHNVVHAIILHQQQQQQQQQQQQQQQPLSQVCFQQSQQQYPSGEGSFQPSQENPQAQGSVQPQQLPQFEEIRNLALETLPAMCNVYIPPYCTIAPVGIFGTNGSGSGGLKMEDVALTLTPGWTQLDSSQVNLYRDEKQENHSSLVSLGGEIQTKSRDLPPVKKLPEKEHGKICHLREDIAQIPTHAEAGEQEGRLQRKQKNAIGS

GDA9_400_2

KTFLILALLAIVATTARIAVRVPVPQLQPQNPSQQQPQEQVPLVQQQQFPGQQQPFPPQQPYPQPQPFPSQQPYLQLQPFPQPELPYPQPQLPYPQPQLPYPQPQPFRPQQPYPQSQPQYSQPQQPISQQQQQQQQQQQQKQQQQQQQQILQQILQQQLIPCRDVVLQQHSIAYGSSQVLQQSTYQLVQQLCCQQLWQIPEQSRCQAIHNVVHAIILHQQQQQQQQQQQQPLSQVSFQQPQQQYPSGQGSFQPSQQNPQAQGSVQPQQLPQFEEIRNLALETLPAMCNVYIPPYCTIAPVGIFGTNGSGSGGGRRCGNGYLEDGEECDCGEEEECNNPCCNASNCTLRPGAECAHGSCCHQCKLLAPGTLCREQARQCDLPEFCTGKSPHCPTNFYQMD g8pw65_400_2

RHQIHELLAPVPGTARKIHSFHFGPEKAVGKIYIQASLHADELPGMLVAWHLKQRLAELEASGHLRHEIVLVPVANPIGLEQVLMDVPLGRYETESGQNFNRRFVDLSEEIGNEIQDLLTDDPQHNLMLIRTSLRDALARQTPGTQLQSLRLALQRLACDADMVLDLHCDFEAVAHLYTTPEAWPQVEPLARYIGAEACLLATDSGGQSFDECFTLLWWQLKERFGERFEIPLGSFSVTVELRGQGDVNHGLASLDCQALIEYLIRFGAIDGEPMPMPELPYPATPLAAVEPVATPVGGLLVYSALPGEYLEAGQLIAQVIDPVNDTVTPVHCRNAGLLYARSLRRMATAGMVIAHVAGTEAYRSGYLLSPGSGSGGPGKVVLVLAGRYSGRKAVIVKV

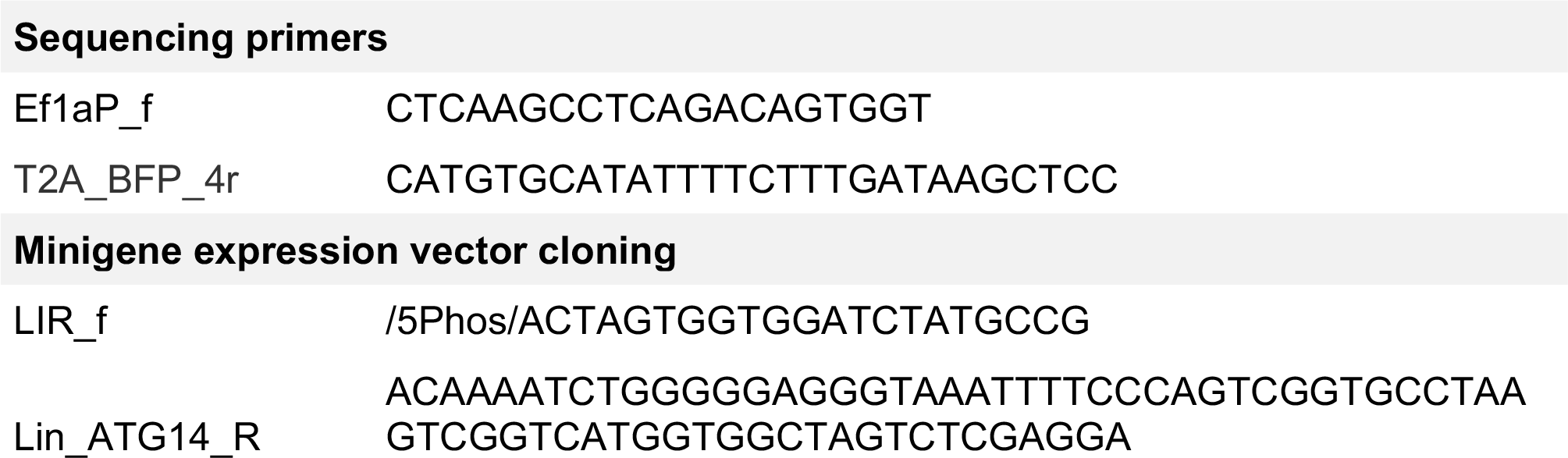
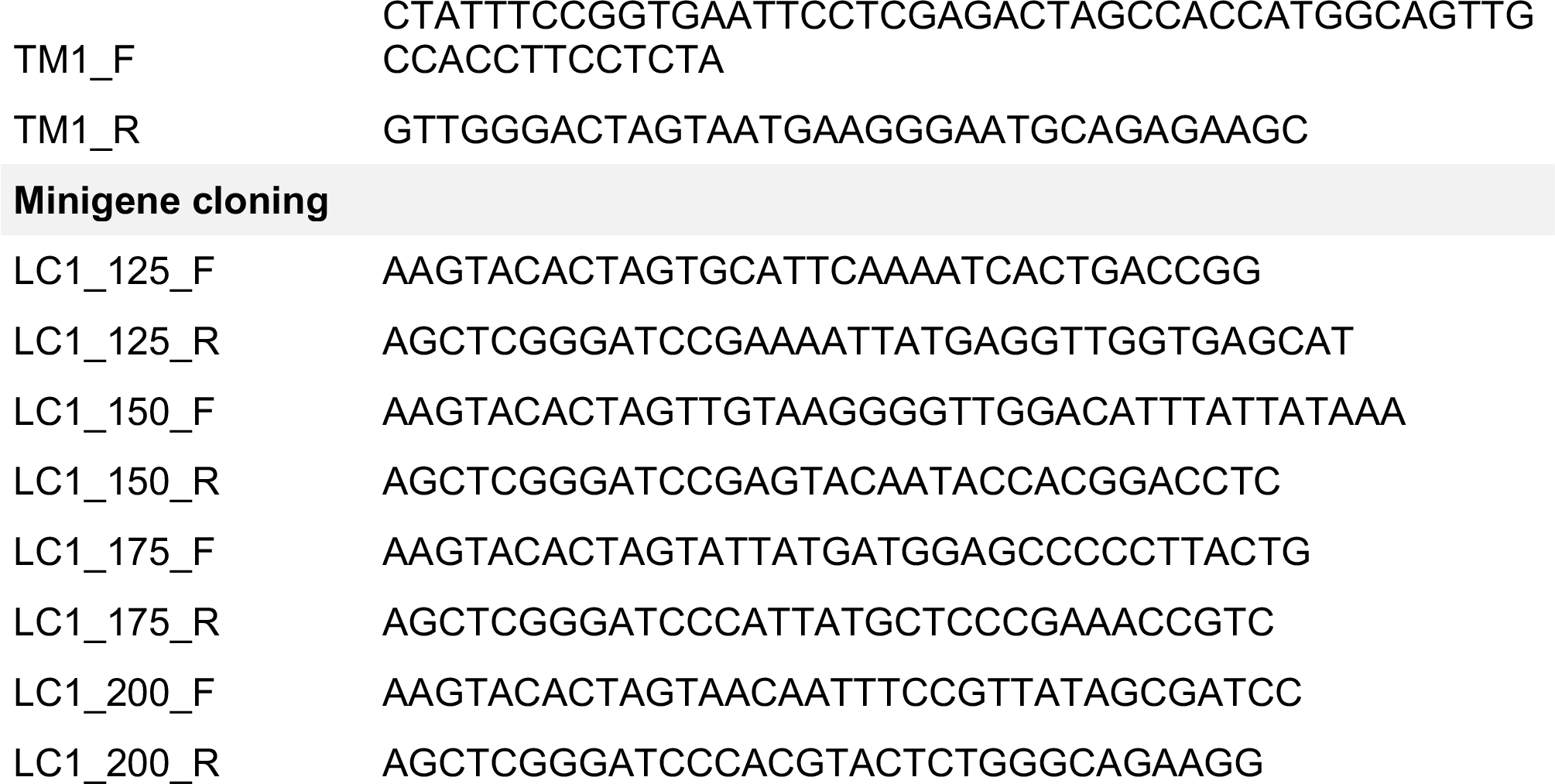
Table of primers used

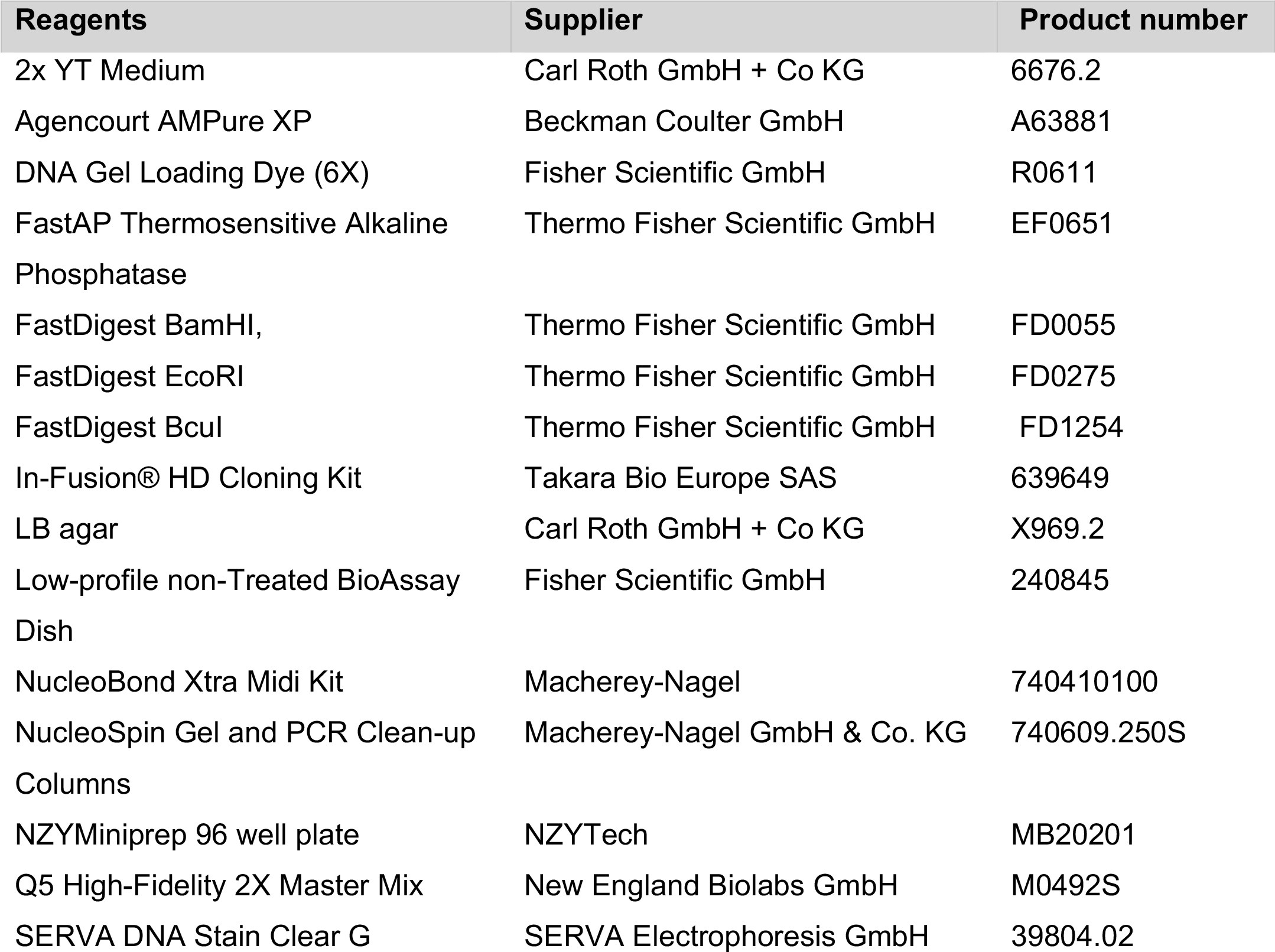
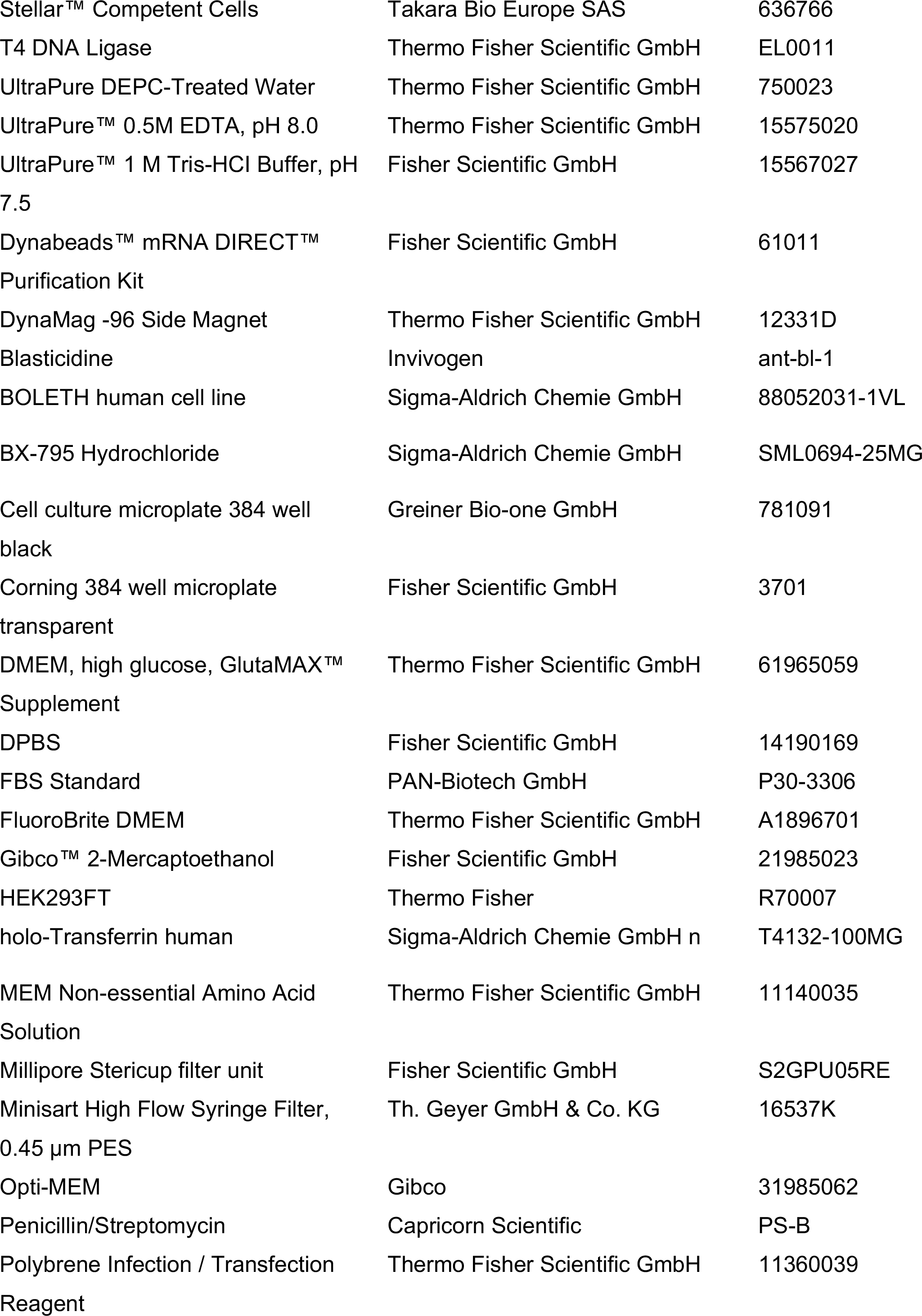
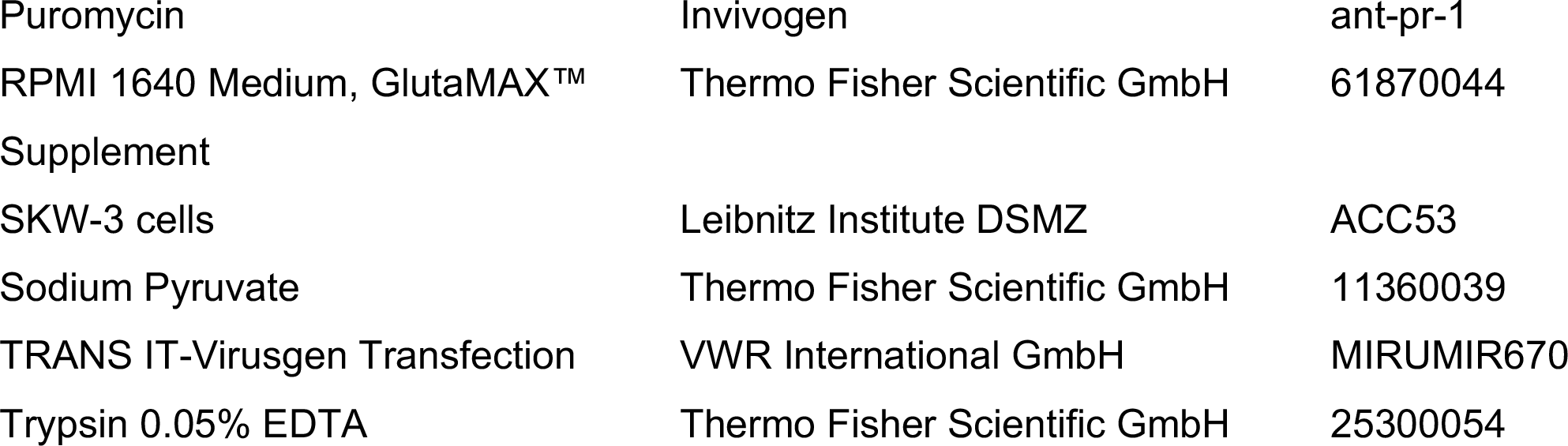
Table of materials used

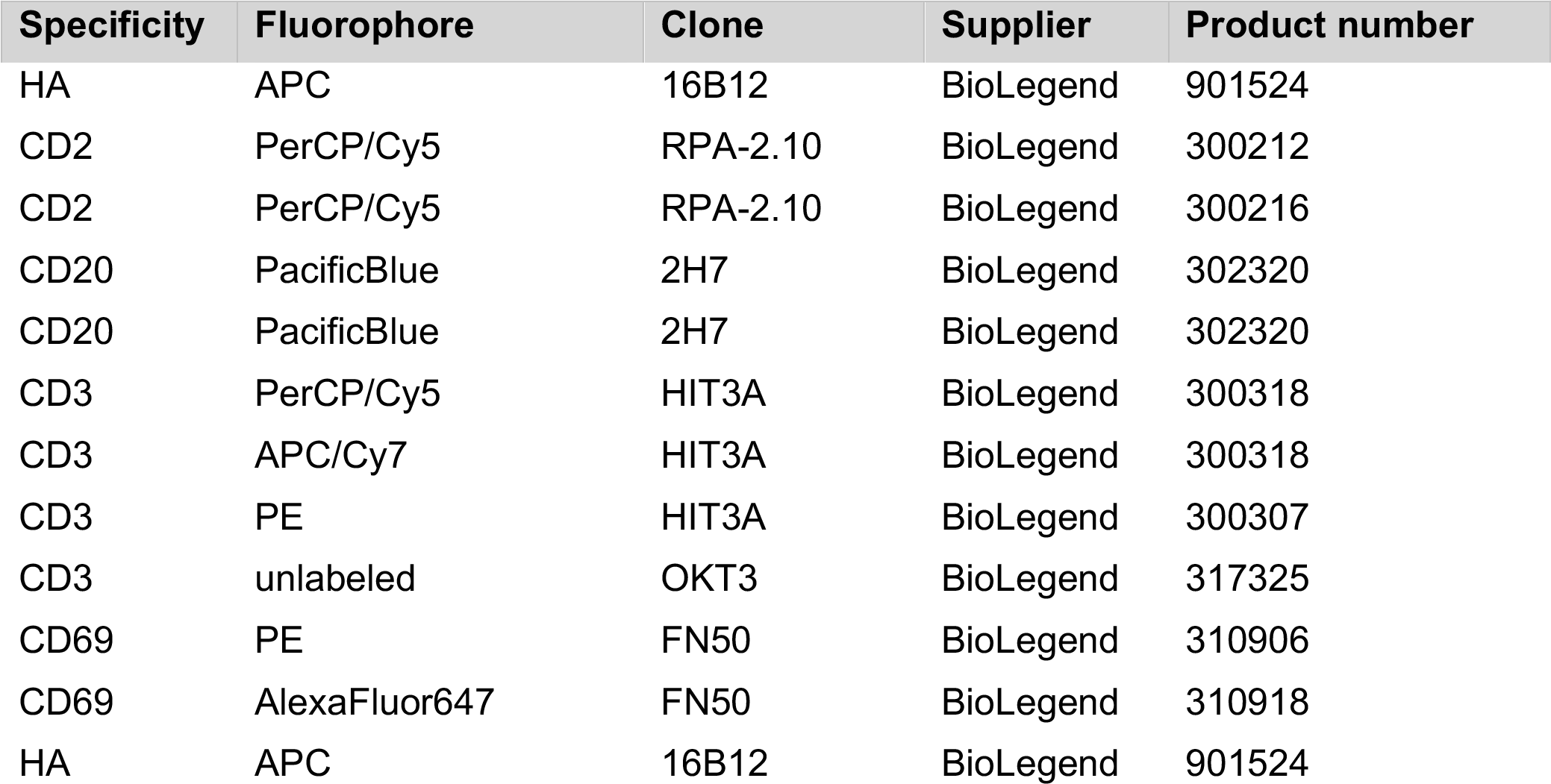
Table of antibodies used

---

**Supplementary Figure 1.**
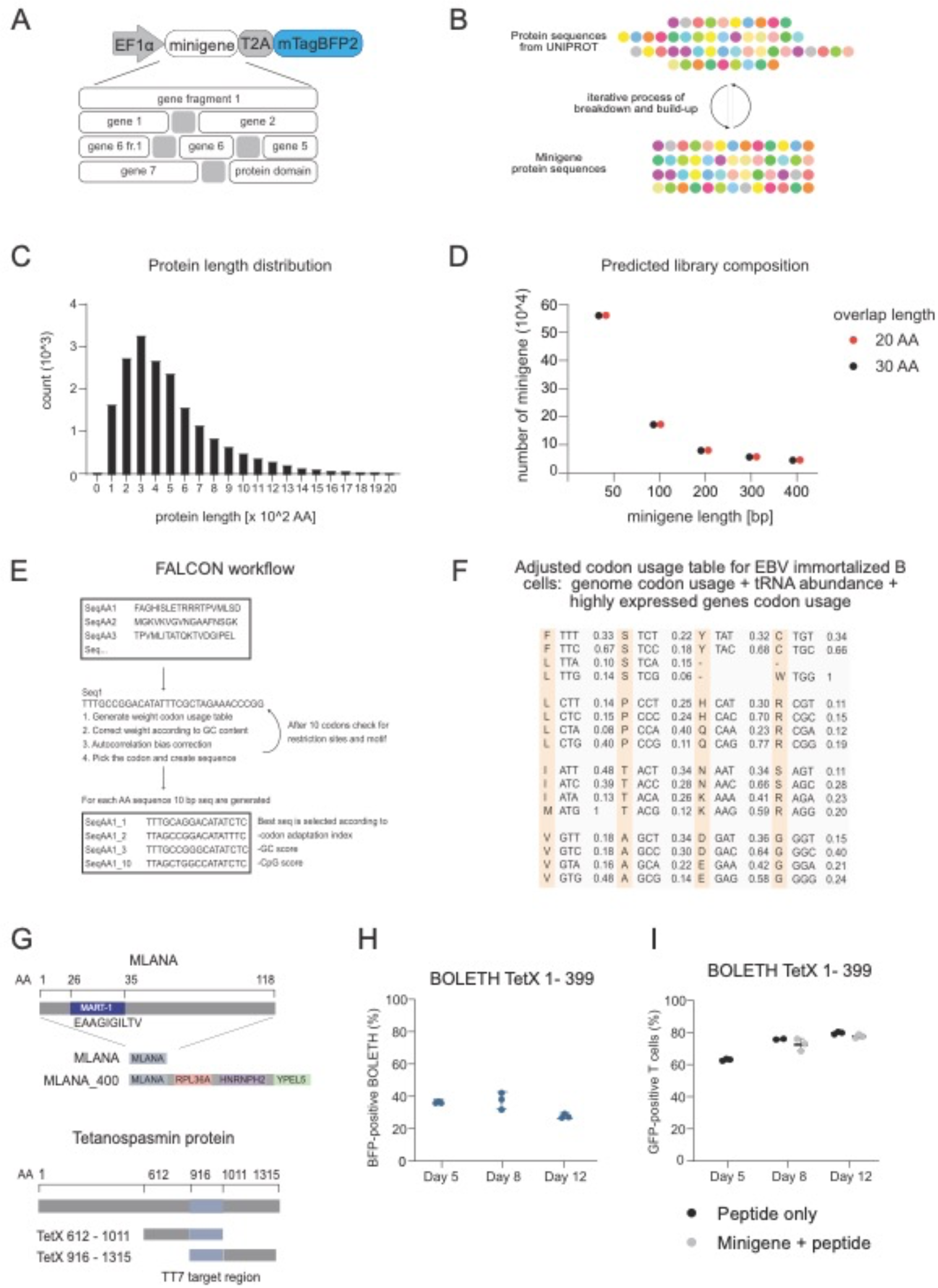
Antigen expression constructs and minigene structure. (A) Schematic representation of the lentiviral vector for expression of an antigen-presenting library, driven by an EF1α promoter. The ORF contains a minigene linked to mTagBFP2 with a T2A peptide. Minigene sequences can be composed of a gene fragment, multiple short genes linked together, genes linked with gene fragments, or a full-length gene linked to a “space-filling” protein domain. (B) Schematic representation of the iterative process for minigene design. Sequences are downloaded from e.g. UniProt, go through iterative process of break down and build-up until all the sequences reach the wanted length. (C) Analysis of protein lengths across the human proteome (D) Predicted number of elements in a proteome-wide minigene library considering minigene length and tiling density (20 aa overlap in black and 30 aa overlap in red.) (E) Schematic representation of the FALCON backtranslation process. (F) Adjusted codon usage table for EBV-immortalized B cells with frequencies for each codon. Genome codon usage, tRNA abundance, and highly expressed gene codon usage are considered. (G) Upper, MLANA protein containing the MART-1 peptide compared to the MLANA_400 minigene, a 400-AA composite into which the full-length MLANA ORF was inserted. Lower, schematic coordinates of the *C. tetani* tetanospasmin protein, which contains the target region (highlighted in blue) for the TT7 TCR. The TetX 612 - 1011 minigene contains the target region at the C-terminal region, while the TetX 916 - 1315 minigene contains the target region at the N-terminal region. (H) mTagBFP2 expression of the transduced BOLETH cells used at each time point. (I) Class II HLA antigen loading control experiment; grey circles, T-cell activation using minigene-transgenic BOLETH cells supplemented with pulsed peptides; black circles, co-culture with freshly peptide-pulsed B-LCLs.

**Supplementary Figure 2.**
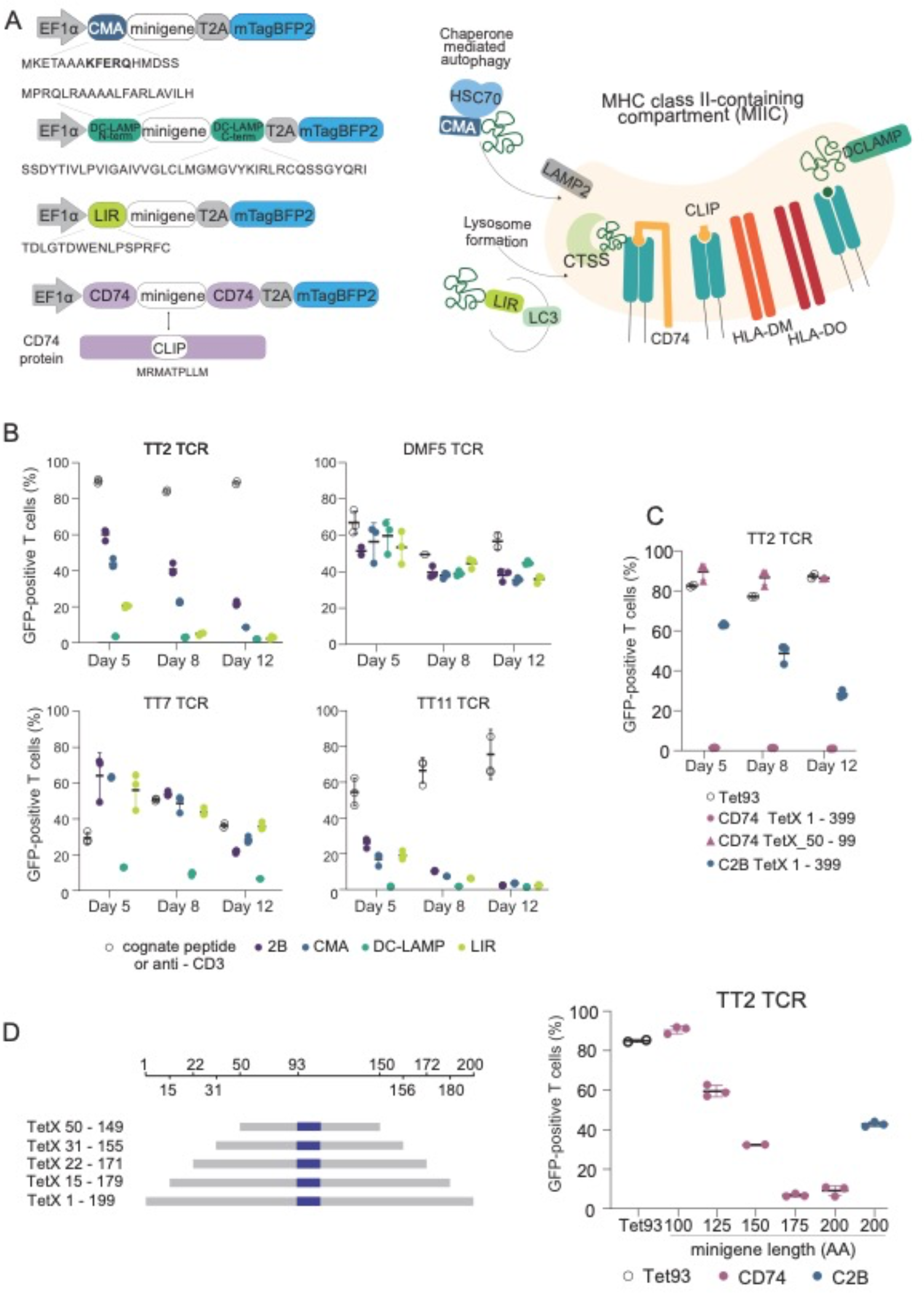
Lysosome localization strategies and CD74 fusion for class II minigene processing and presentation. (A) Schematic of class II processing constructs (left) used in this study and their projected pathway entry points (right). (B) Flow cytometric analysis of co-cultures at day 5, day 8, and day 12 after transduction with CMA, DC-LAMP, and LIR constructs for minigene expression. TCR:minigene pairs used: TT2:TetX 1 - 399, TT11:TetX 612 – 1011, TT7:TetX 612 – 1011, DMF5 – MLANA^A27L^. 1 μM Tet93, 1 μM Tet612, 2.5 μg/ml of α-CD3, and 10 μM MART-1, respectively, were used as positive controls for reporter cell activation, expressed as the percentage of GFP positive T cells following co-culture. (C) Reporter cell activation at days 5, 8, and 12 following transduction with TetX 50 – 99 or TetX 1 – 399 minigenes, both cloned in the 2B or CD74 backbones. Reporter T cells expressed the TT2 TCR; B cells loaded with 1 μM Tet93 peptide served as a control for the cognate interaction. (D) Left, schematic representation of the minigenes used, aligned to their respective positions in the tetanospasmin protein. Colored boxes indicate the TT2 peptide epitope. Right, co-culture results at day 5 using minigenes of increasing length cloned in CD74 or C2B construct. 1 μM of Tet93 peptide served as the positive control.

**Supplementary Figure 3.**
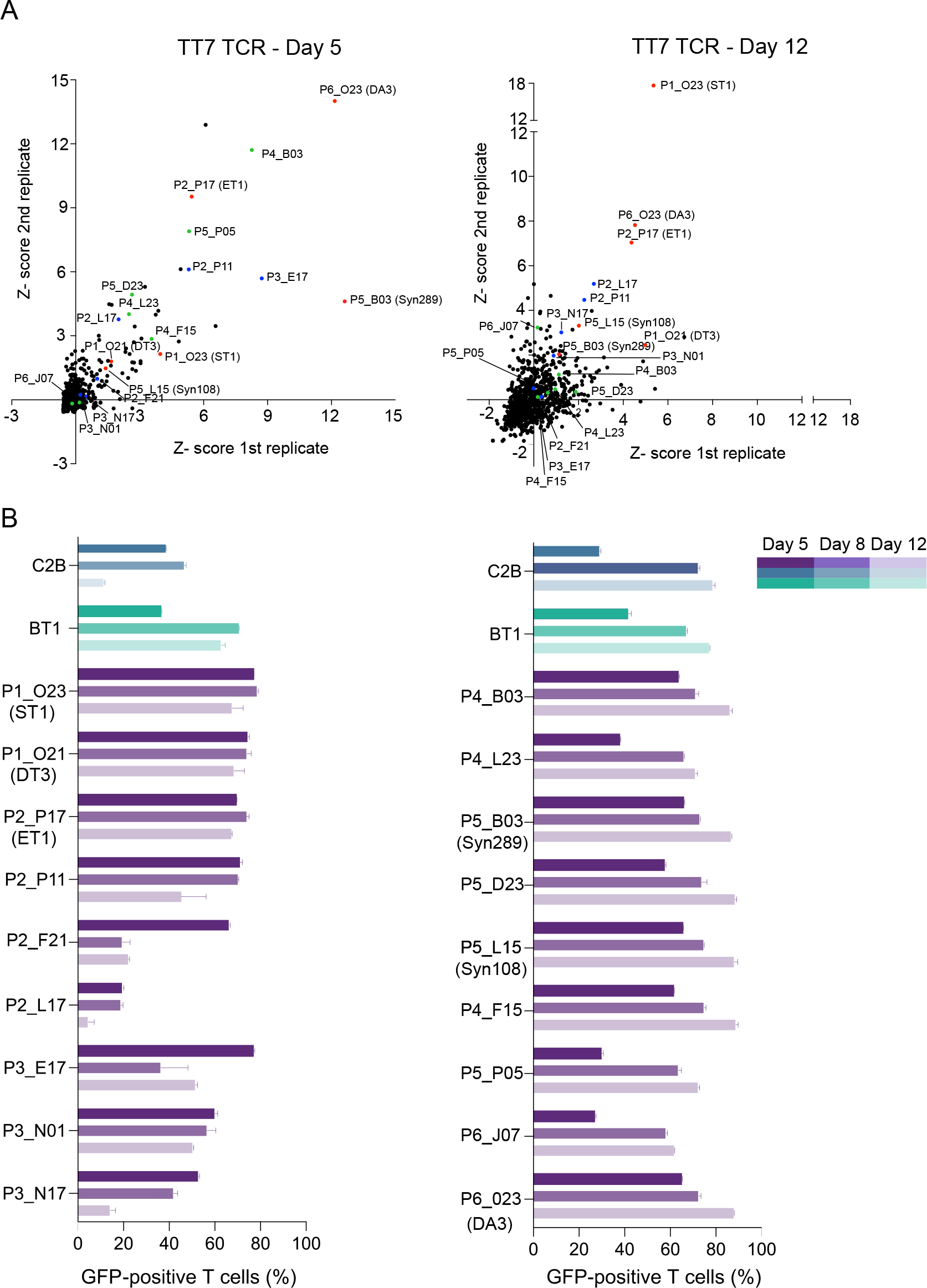
TM_ER domain library functional screening. (A) Results of a plate-based screen of TM_ER domain library elements. Top panels show each of the 1152 colonies screened. Each dot represents the replicate value Z-scores of GFP median fluorescence for a given clone in co-culture with a cognate TCR:minigene pair. The left plot depicts results at day 5 and the right plot depicts results at day 12 post-transduction. Blue circles indicate sequences also identified in the first rounds of screening; green circles are those from additional screening rounds. Red circles indicate clones tested with the TT2, TT11, and TT7 TCRs in Figure 3. (B) Results of two rounds of validation assays, on the left the first round (blue circles above) and on the right the second (green circles). GFP expression levels (% positive cells) were recorded at day 5, 8, and 12 in comparison with the 2B and BT1 constructs.

**Supplementary Table 1.**
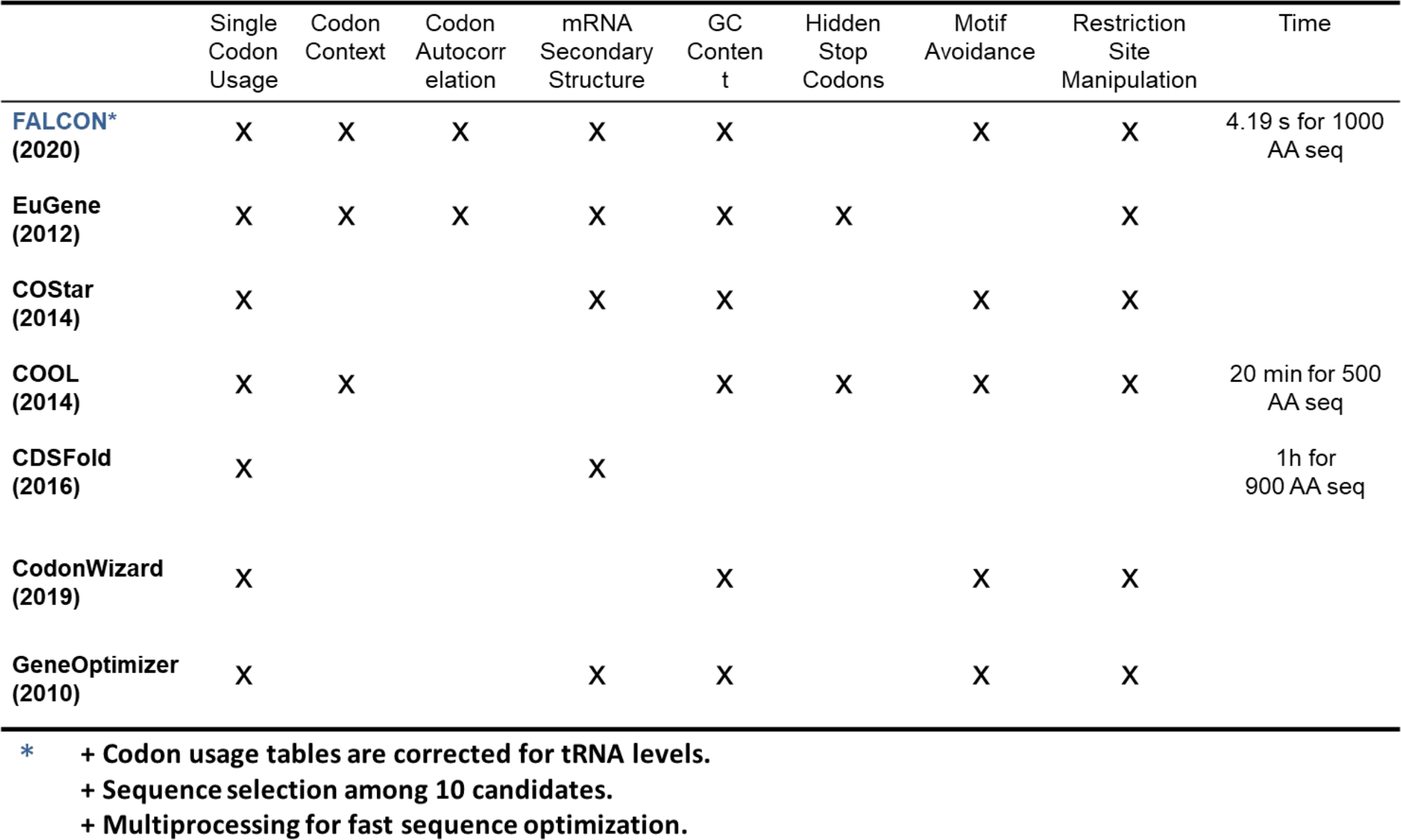
FALCON back-translation script features.

**Supplementary Table 2.**
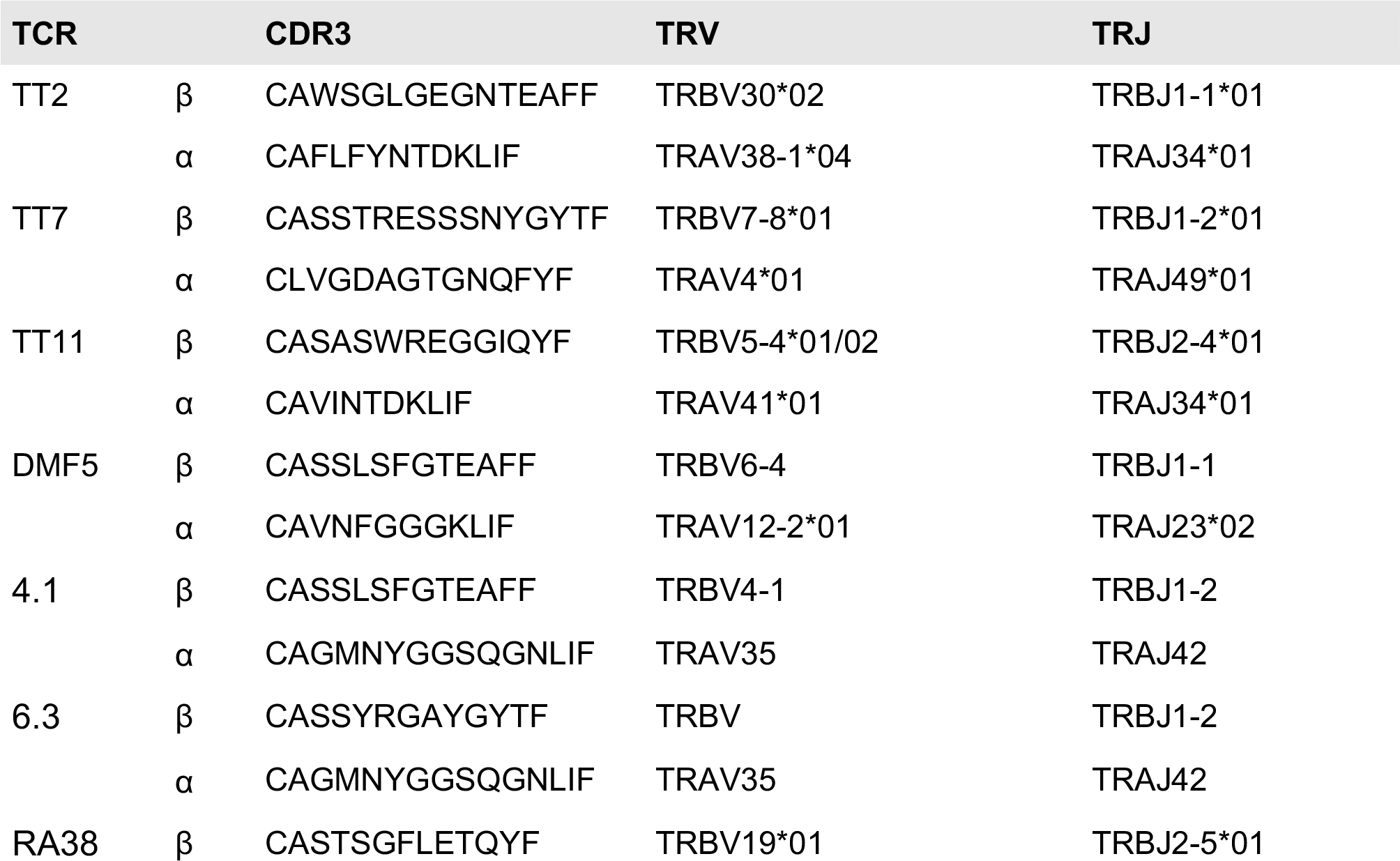
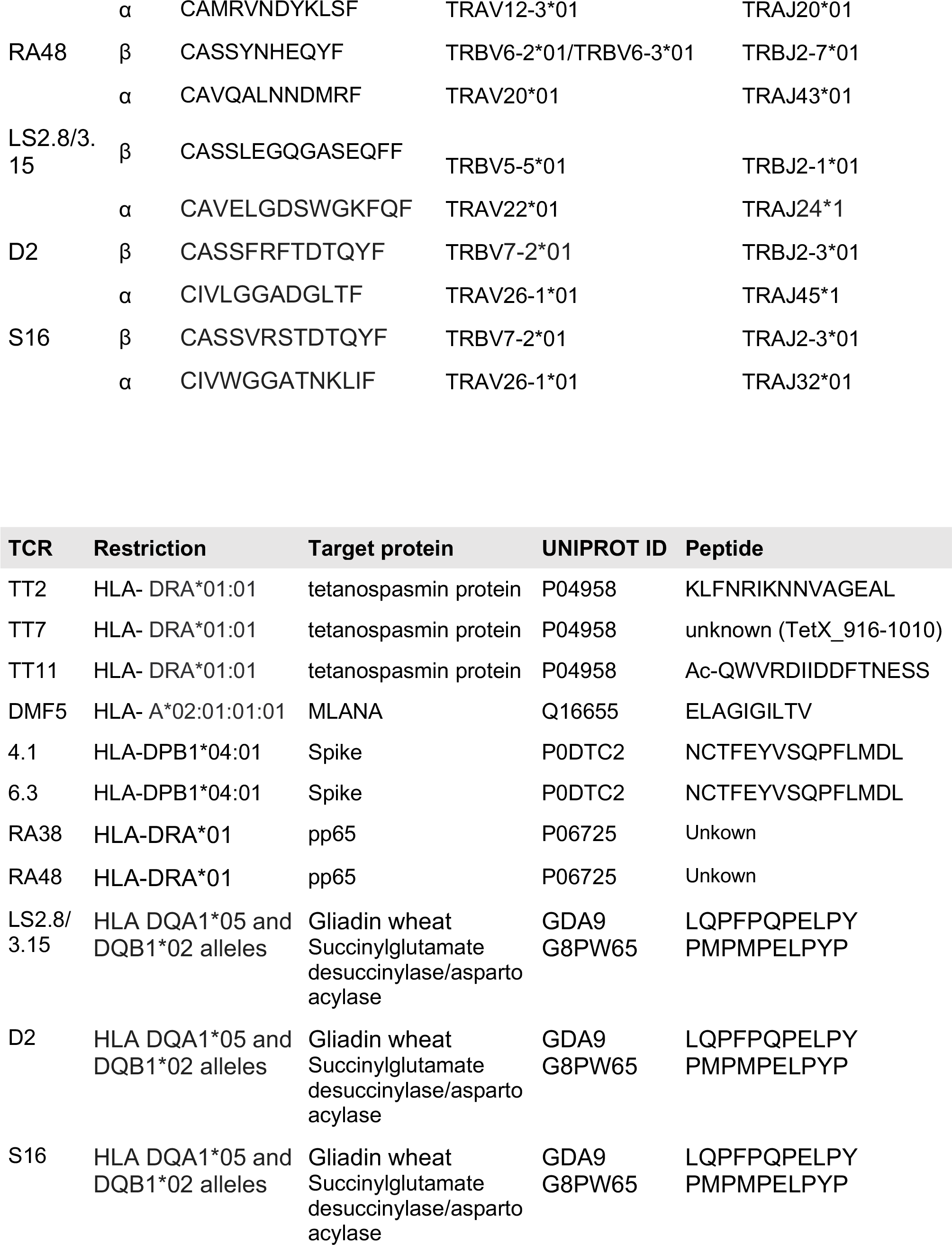
Table of TCRs.

